# Modulating the unfolded protein response: Impacts of radiation on the response of prostate cancer cells to ONC201

**DOI:** 10.1101/710400

**Authors:** Francesca Amoroso, Kimberley Glass, Francisco Liberal, Rebecca E. Steele, Sarah Maguire, Rohinton Tarapore, Joshua E. Allen, Sandra Van Schaeybroeck, Karl T. Butterworth, Kevin Prise, Joe M. O’Sullivan, Suneil Jain, David J. Waugh, Ian G. Mills

**Affiliations:** Centre for Cancer Research and Cell Biology, Queen’s University Belfast, United Kingdom, BT9 7AE; Faculdade de Ciências e Tecnologia, Universidade Nova de Lisboa, Portugal, 2825-516; Breast Cancer Now Toby Robins Research Centre, The Institute of Cancer Research, London, United Kingdom, SW3 6JB; Research & Development, Oncoceutics Inc., Philadelphia, PA 19104, USA; The Northern Ireland Cancer Centre, Belfast City Hospital, United Kingdom, BT9 7AB; Queensland University of Technology, Brisbane City QLD 4000, Australia; Nuffield Department of Surgical Sciences, University of Oxford, United Kingdom, OX3 9 DU

## Abstract

Prostate cancer (PCa) is the most common non-cutaneous cancer in men and a notable cause of cancer mortality when it metastasises. Localised disease is mostly treated with surgery or radiotherapy. As PCa develops and treatment resistance emerges, the unfolded protein response (UPR) arises as an important adaptive biology co-amplifying with key cancer drivers [1]. The UPR can be cytoprotective but when acutely activated can lead to cell death. In this study we sought to enhance the acute activation of the UPR using radiation and ONC201, previously reported to be an UPR activator [2]. We found that treating PCa cells with ONC201 quickly increases the expression of components in all arms of the UPR – ATF4, ATF6 and IRE1-XBP1 – culminating in the subsequent cell death. During this time window between UPR activation and cell death we tested the priming effect of short-term administration of ONC201 on radiation responses. Pre-treatment with ONC201 for 24 hours prior to irradiation led to enhanced cytotoxicity compared to radiation alone assessed by cell viability and clonogenic assays. With priming, RNA-Seq analysis showed a sustained suppression of transcripts encoding cell cycle regulators as well as components of the DNA damage response pathways. Phenotypically this was reflected in enhanced cell cycle arrest and induction of necrosis and apoptosis. Furthermore, we demonstrated that short-term administration of inhibitors of cell cycle regulators (Dinaciclib and BI2536), could replicate this priming effect. Thus, we propose future studies to assess the impact of the short-term administration of drugs targeting the UPR and cell cycle regulation to enhance radiotherapy response.

## Introduction

Prostate cancer (PCa) is the most common cancer diagnosed in men and the second most common cause of cancer death after lung cancer. According to recent projections, prostate cancer incidence rates are predict to rise by 12% in the UK between 2014 and 2035, to 233 cases per 100,000 males by 2035 [3]. Clinically localised PCa is treated using radical prostatectomy or radiotherapy to remove or destroy the cancer cells confined within the prostate capsule. However, 10 to 15 % of the patients are diagnosed after their cancer has spread and present with advanced or inoperable disease [4].

The prostate is a specialized accessory gland with a high secretory capacity. During cancer progression, cells experience mitogenic pressure and intracellular stress (e.g., metabolic pressure to rapidly grow and divide), detected by the endoplasmic reticulum (ER) as an accumulation of misfolded proteins. When the cells are not able to cope with the overload, the unfolded proteins accumulated in the ER, trigger an adaptive response called the Unfolded Protein Response (UPR) [5]. Attempting to clear the unfolded proteins and increase the capacity of the ER, the UPR activates several molecular pathways. Here, the so-called ER stress sensors PERK, IRE1 and ATF6 play a central role in the initiation and regulation of the UPR [6] [7] [8] [9]. Previously, several studies have reported the activation of the UPR during tumour transformation and progression, leading to the acquisition of adaptive phenotypes to restricted nutrient supplies and therapies [10]. Upstream elements like XBP1 and ATF6 are upregulated in hepatocellular carcinomas [11], in a range of breast cancer cell lines [12], colon cancer and melanoma [10]. Although the UPR is generally viewed as a cytoprotective response, prolonged ER stress can directly regulate the cell death machinery through the activation of CHOP [13] [14]. One of the mechanisms by which CHOP promotes apoptosis involves its ability to decrease anti-apoptotic Bcl-2 levels and stimulate the release of cytochrome C into the mitochondrial lumen to result in the activation of apoptotic caspase 3 [15]. Being a major secretory organ, the prostate is particularly reliant on the proper functioning of the ER and is vulnerable to agents or conditions that cause ER stress. Several studies have pointed to a positive association between ER/UPR markers and the development of prostate cancer [1]. Even though the activation of the UPR is known to be the result of the accumulation of unfolded proteins in the ER [1], treatments such as radiotherapy also increase stress and perturb general cellular homeostasis.

Radiotherapy causes double-strand DNA damage [16] [17] [18]. If unresolved, the radiation-induced DNA damage can lead to the production and the accumulation of unfolded and/or misfolded proteins in the ER [19]. Recent literature has shown that radiation exposure of glioblastoma stem cells activates key components of the UPR culminating in autophagosome formation [20]. Moreover, the overexpression of UPR genes encoding GRP78 and GRP94 has been extensively associated with radio-resistance in multiple cancer types, including breast, pancreatic and gastric cancers [21] [22, 23]. Recently Drake and co-workers demonstrated that therapeutic doses of radiotherapy led to an upregulation of GRP78 in the 72% of colorectal cancer cases receiving treatment [24]. Targeting the UPR may provide an opportunity to enhance responses to radiotherapy.

ONC201 is an inhibitor of the dopamine receptors DRD2/3 and has previously been reported to induce apoptosis in haematological malignancies and solid tumours [2]. Subsequently it was shown that apoptosis in response to ONC201 treatment was triggered by the activation of the ATF4 axis of the UPR [25]. ONC201 has been recently shown to be effective in early stage clinical trials in a number of cancer types [26] [27]. In prostate cancer activation of the IRE1-XBP1 axis a the UPR contributes to tumorigenesis in contexts in which the driver is the androgen receptor [28]. In prostate cancers characterised by c-Myc overexpression and PTEN mutations, the ATF4 axis of the UPR plays a pro-survival role [29]. Here, we assess the impact of the ONC201 on all axes of the UPR to determine whether it can be used to enhance radiation response in PCa models.

## Results

### CHOP expression influences radiation response

The activation of the UPR in known to be the result of the accumulation of unfolded proteins in the ER [1]. CHOP is known to be a pro-apoptotic transcription factor expressed in response to acute stress which can include therapeutic stress such as genotoxic stress to irradiation of cancer cells. To confirm this, we generated stable knock downs of (CHOP) in PC3 cells and irradiated both the knockdown clone and the scrambled control with single doses of radiation (**Figure 1, a and b**). Cells were up to 20% more resistant at 72hours to the highest dose (8Gy – white bars vs black bars). Moreover, without radiation, PC3-shCHOP grew faster than the shScram (if we compare white bars vs black No Rad). This confirmed that CHOP expression contributes to radiation responses and prostate cancer cell survival.

**Figure 1.**
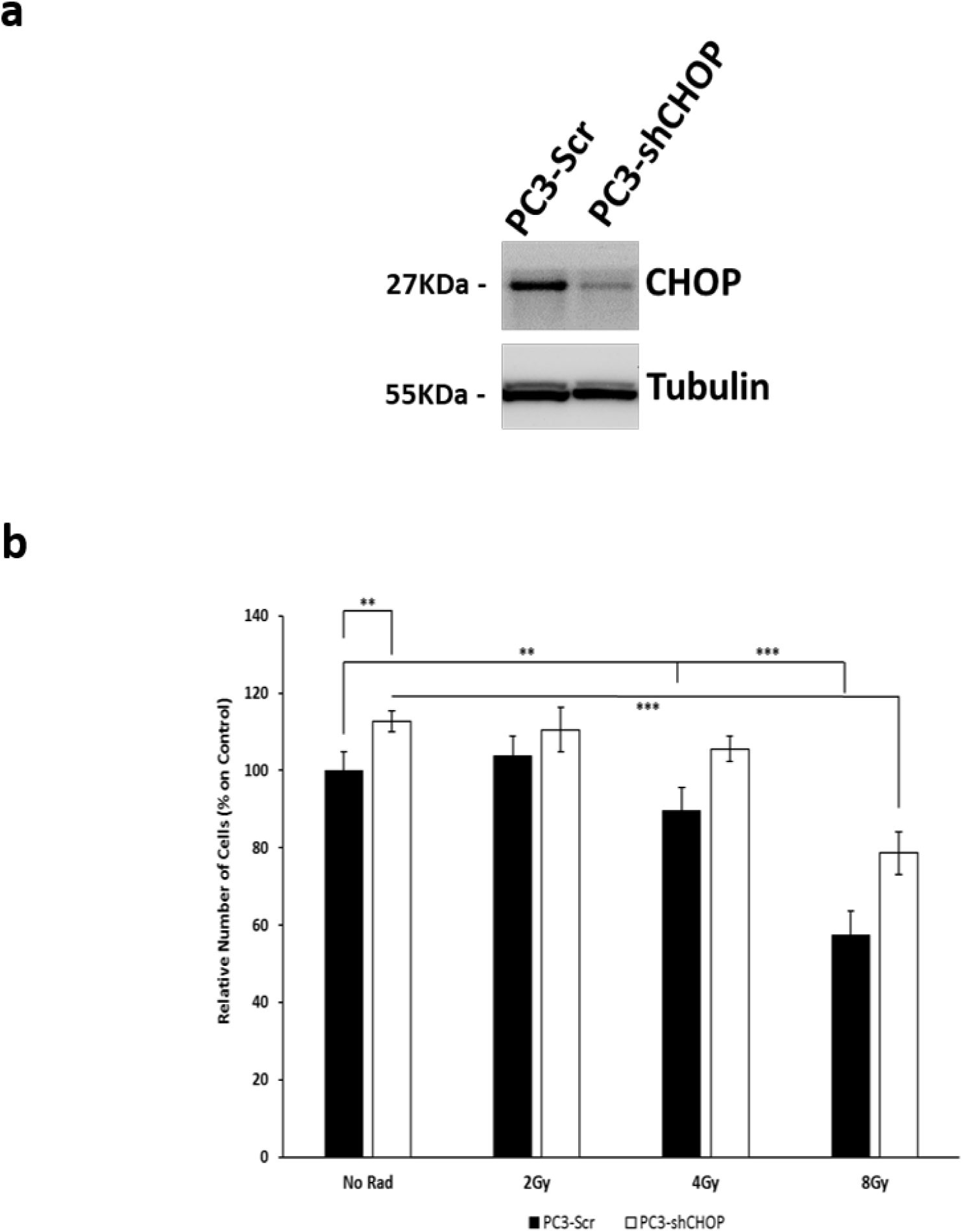
**a)** Expression of CHOP in PC3 cells Scramble (line 1) and silenced through shRNA infection (line 2). **b)** Effects of radiation (2Gy, 4Gy and 8Gy) on cell counts of PC3-Scr and PC3-shCHOP. Cells have been counted at 72hours from radiation and are expressed as normalised on the relative cell counts at T0 (narrowed bar). Knocking down the expression of CHOP makes PC3 cells less sensitive to radiation. One way ANOVA test has been run for statistical significance, *P<0.05, **P<0.03, ***P <0.001 (±SD).

### Chronic activation of the Unfolded Protein Response with imipridone ONC201 induces cell death

Given that downregulation of CHOP led to cell survival, we hypothesized that the activation of pro-apoptotic arms of the UPR may enhance cell death. To achieve this, we utilized two different compounds (HA15 and ONC201) known to activate the UPR in other models [2, 30]. Of the two, only the administration of ONC201 (5, 10 and 15 µM, [31]) significantly induced cell death at 72hours (**Figure 2, a**), whereas HA15 showed a cytostatic effect at all the concentrations tested (1, 2.5 and 5 µM, [30]) (**Supplementary Fig. 1, a**). Interestingly, the cytotoxic effect observed with ONC201 at 72hours arose after significant and dose-dependent increases in the expression of UPR components at 24hours (**Figure 2, b**) and the activation of Caspase 3 at 48hours (**Figure 2, c**). At 72 hours we then observed a significant and dose-dependent increase in apoptosis (blue and red bars) and necrosis (**Figure 2, d**). Cell Cycle analysis at 72hours after the administration of ONC201 (**Figure 2, e**) showed a significant and dose-dependent increase of up to 30% in the proportion of cells in S phase and a significant increase up to 20% in the proportion of cells in the G2/M phase. There is evidence that ONC201 modulates the cell cycle in haematological malignancies and leukemic stem/progenitor cells [32]. In this setting ONC201 causes p53-independent apoptosis and a delayed S phase of the cell cycle [32].

**Figure 2.**
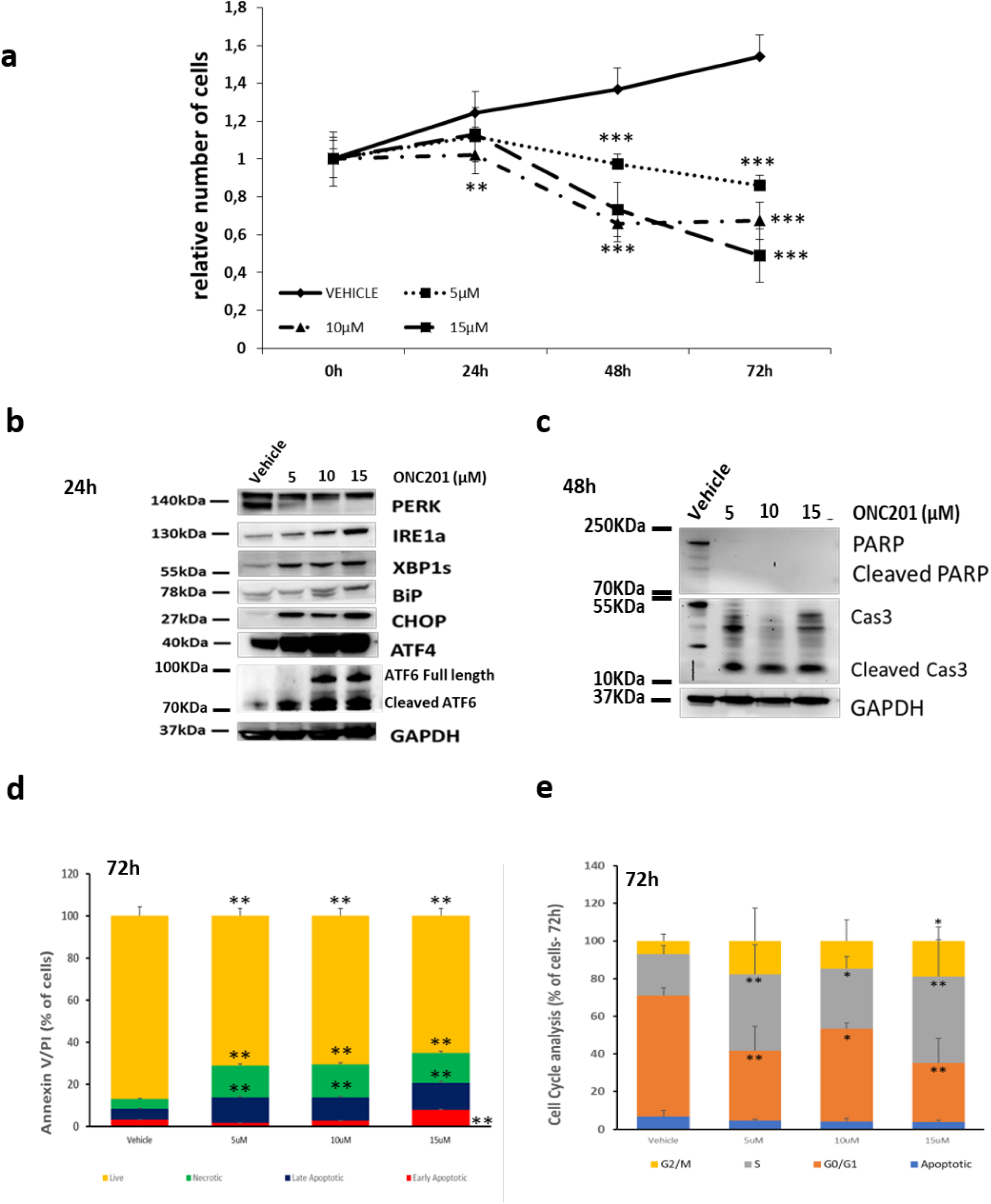
**a)** ONC201 has a cytotoxic effect on PC3 cells treated up to 72h. **b)** Expression of proteins composing the UPR in PC3 cells treated with Vehicle and ONC201 (5µM, 10µM and 15µM for 24 hours. All the proteins of the UPR are overexpressed after 24hours of treatment with ONC201. **c)** ONC201 induces cell death trough the cleavage of Caspase3 at 48 hours. **d)** ONC201 induces cell death by both apoptosis and necrosis. **e)** ONC201 Induces cells to exit G0/G1 phase of the Cell Cycle to enter the S phase. One way ANOVA test has been run comparing treated vs Vehicle, *P<0.05, **P<0.03, *** P <0.001 (±SD).

### Priming Prostate Cancer cells with ONC201 increases the efficacy of irradiation

Having observed a clear temporal progression from UPR activation at 24hours to cell death at 72hours we sought to test whether short-term administration of ONC201 could prime subsequent cell death responses to irradiation (**Figure 3, a**). We used 3 doses of ONC201 (5, 10 and 15 µM) and 3 doses of radiation (2, 4 and 8Gy) (**Figure 3, b**). After 24hours the compound was washed out and the cells were irradiated. We observed 30-50% more cell death by pre-treating the cells with ONC201 than we observed with radiation alone. We therefore calculated the Radiation Enhancement Ratio (RER) at 72hours from radiation (**Figure 3, b**, right side). The RER gave us a measurement of the radio-sensitizing effect that priming with ONC201 had on the PC3. A 4Gy dose of radiation was twice as effective when following on from pre-treatment with with ONC201 (5 and 10µM) compared to radiation alone. We confirmed that pre-treatment with ONC201 (5 and 10µM) was indeed synergistic with radiation (**Figure 3, c**). By contrast the highest pre-treatment dose of ONC201 (15µM) was not synergistic (data not shown) and this may reflect the cytotoxicity of the drug at this dose as a single agent. The clonogenicity of PC3 cells was also dramatically reduced by ONC201 pre-treatment (**Supplementary Fig. 1, b**). Blotting cell lysates we found that the priming effect of ONC201 led to sustained increases in the expression of proteins in all arms of the UPR at 72hours and particularly of IRE1, ATF4 and CHOP but also a reduction in XBP1s (**Figure 3, d; Supplementary Fig. 1, d and Supplementary Fig. 2, a**). At 72hours, more intriguingly, we observed the upregulation of distinct components of the UPR dependent on the treatment. OMC201 treatment led to increases in PERK, ATF4 and CHOP whereas radiation led to increases in IRE1a and ATF6. (**Figure 3, d – left side**). Priming prostate cancer cells with ONC201 for 24hours followed by radiation, maintained the over-expression of both signatures at 72hours (PERK/ATF4/CHOP plus IRE1a/ATF6) (**Figure 3, d – right side**). At an earlier time point, 24hours post-irradiation, pre-treatment with ONC201 by contrast led to a significant increase solely in CHOP levels (**Supplementary Fig. 1, d and e**). Given that the treatment with ONC201 followed by radiation was inducing the overexpression of proteins of the UPR known to promote cell death, we sought to determine how the cells were dying. Once again, we observed very distinct responses when cells were treated with radiation alone or ONC201. Radiation induced cell death primarily by apoptosis (ca. 10% of the cells) and by contrast ONC201 induced necrotic cell death in up to the 20% of the cells. The overall effect of the sequential treatment on cell survival was the activation of both necrosis (green, up to 30%) and apoptosis (red and blue, up to 20% of the total population), combining the cell death characteristics of each individual treatment (**Figure 3, e**).

**Figure 3.**
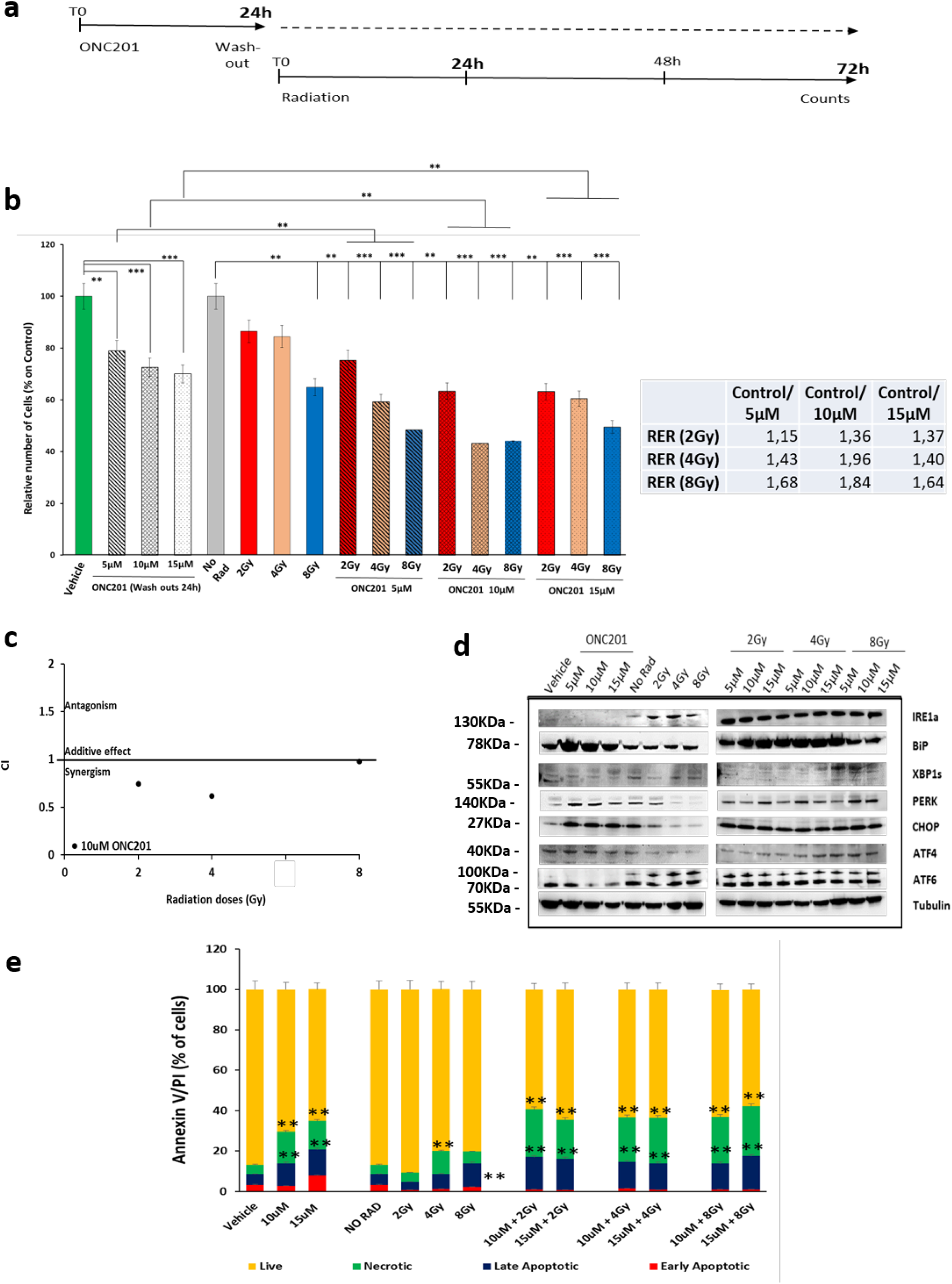
**a)** Schematic representation of the treatment in which PC3 cells have been primed with ONC201 before being radiated with single doses of Xrad. **b)** Priming PC3 cells to radiation (Xrad) with ONC201 for 24hours impacts cells survival at 72hours. Radiation Enhancement Ratio (RER) is shown and calculated out of 3 independent experiments. **c)** ONC201 10µM has a synergistic effect with radiation (CompuSyn). **d)** Representative of n=3 WB analysis of the expression of all the components of the UPR upon priming PC3 cells to radiation (Xrad) with ONC201 for 24hours. Samples were harvested at 72hours from the last radiation. Densitometric analysis of n=3 WB are shown in Supplementary Fig.2, (a). **e)** Cell death analysis through Annaxin V/PI detection of PC3 treated with ONC201 (5 and 10µM) and Radiation (as described in (a). Samples were harvested at 72h. One way ANOVA test has been run, *P<0.05, **P<0.03, ***P <0.001 (±SD).

### RNA-Seq analysis reveals that ONC201 restricts the expression of cell cycle progression and DNA repair pathway genes

To reveal more about the molecular mechanisms through which ONC201 may function as a radio-sensitizer, we performed RNA-Seq analysis on treated PC3 cells (**Figure 4, a**). Priming PC3 cells to radiation utilizing ONC201 for 24hours (then washed out) and collecting the cells at 24hours and 72hours after irradiation, we observed that our samples clustered according to the timepoints of harvesting (**Supplementary Figure 2, b** – red dots vs blue dots respectively). To identify significant differentially expressed genes we therefore focussed on a single timepoint, 72 hours, post-irradiation and on genes that were significantly overexpressed in surviving cells (**Table 1**). Pathway analysis **Figure 4, b and Supplementary Figure 2, c**) (**Table 2**) revealed enrichment for cells cycle progression and DNA replication and repair pathways. We than generated heatmaps for the genes that were most significantly upregulated in cells surviving radiation (red squares) and downregulated by ONC201 treatment (blue squares) and these included RRM2, MiK67, TYMS, PLK1, CDK1 and CDK2 (**Figure 4, c i**). The RNA-Seq data also confirmed that ONC201, both in single treatment and as a radiosensitizer, induced increased expression of ATF4 and CHOP at 24hours. In contrast to the increases in the UPR expression we observed at the protein level at 72hours, the transcript levels of factors such as CHOP and ATF4 were downregulated at this timepoint in our RNA-Seq data (**Figure 4, c ii**). The lack of correlation between the expression of UPR transcripts and proteins has been previously reported and may reflect partly an impact on the normal protein folding and turnover functions of cells experiencing stress [33] [34]. This may reflect feedback compensation for post-transcriptional defects. We validated transcripts encoding RRM2, CDK2, CDK1, and PLK1 based on their significant differential expression and regulatory roles in the control of cell cycle progression and DNA damage repair (as shown in **Figure 4, e**) (**Figure 4, d**). Three of these transcripts encode proteins that are known to be druggable, PLK1, CDK1 and CDK2 and are key cell cycle regulators. We further investigated the impact of ONC201 on the expression of these factors and confirmed by Western blotting that polo-like kinase 1 (PLK1) was downregulated at the protein level by ONC201 despite being overexpressed when cells were treated with radiation alone (**Supplementary Figure 4, a**). We also validated the expression of MiK67 gene because coding for a protein (Ki-67) commonly used as a marker of proliferation, thus corroborating the phenotypic data we described previously.

**Figure 4.**
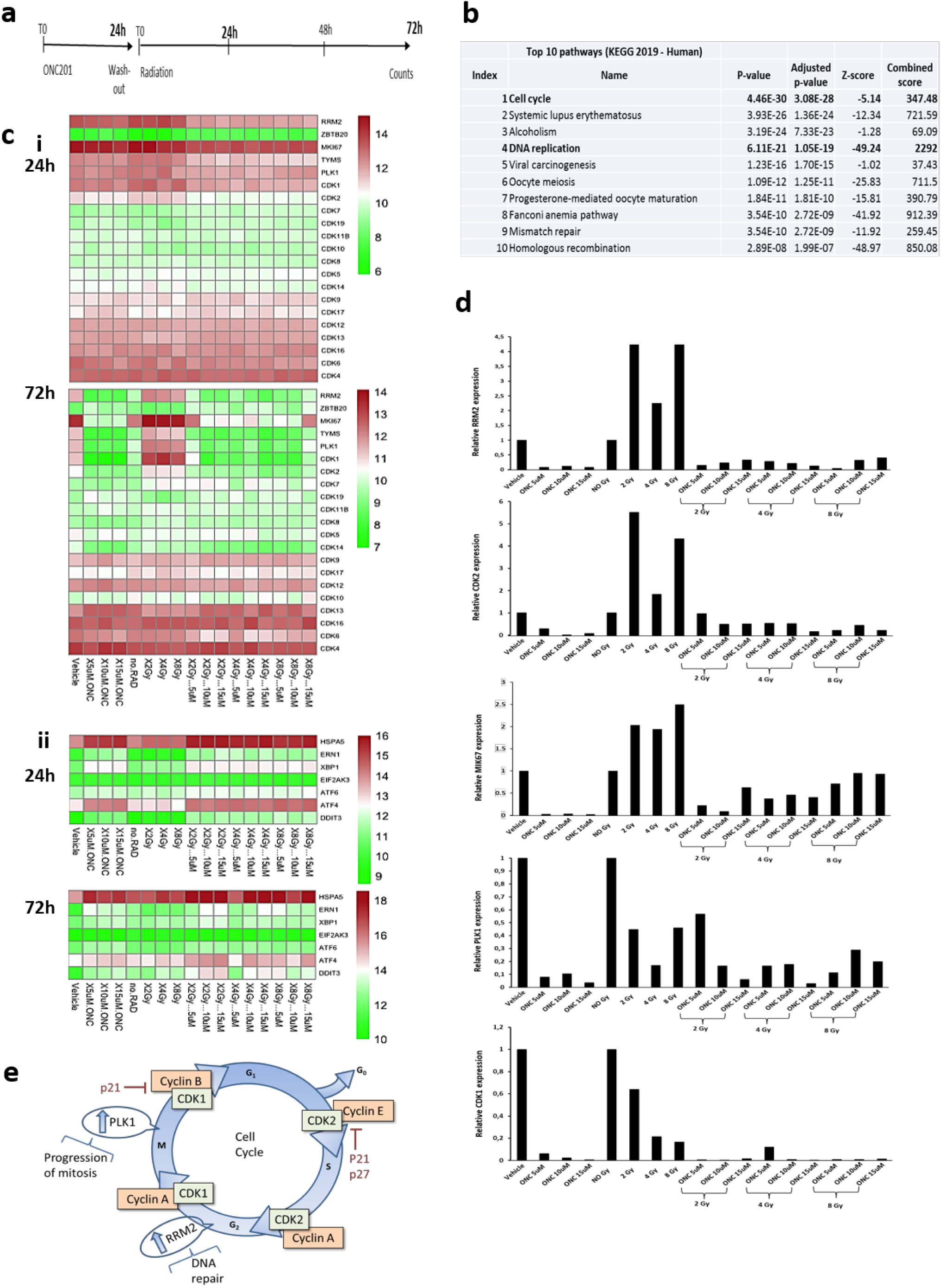
**a)** Schematic representation of the treatment in which PC3 cells have been primed with ONC201 before being radiated with single doses of Xrad. **b)** Pathway analysis of our genes of interest in KEGG Human 2019 pathway database reveals that the most impacted pathway is cell cycle related (iEnrichr web tool – [60] [61]). **c)** Heatmaps analysis of RNAseq data (log2) shown in (a). Genes listed are differentially regulated at 24h and 72h from the final radiation and cell cycle related. **d)** Validation though RT-PCR of differentially expressed genes at 72hours from the final radiation dose (*RRPM2, CDK2, PLK1, MIK67, CDK1). **e)** Schematic representation of the Cell Cycle check points (*RRPM2, CDK2, PLK1, MIK67, CDK1).

### ONC201 induces an expansion of the S phase of the cell cycle

To further investigate the role of ONC201 in modulating genes controlling the cell cycle progression and the DNA damage response, we performed cell cycle analysis on cells treated and collected at 72hours, as summarized in Figure 4, A and 5, B (**Figure 5, a**). We observed a doubling of the S phase population and an expansion of the G2/M phase population (up to 20% more) upon treatment with ONC201 alone. These increases were even greater when the radiation response was primed by ONC201 pre-treatment. Thus, we hypothesised that the impact of ONC201 as a radio-sensitizer was achieved both by activating the pro-death branches of the UPR and restricting cell cycle progression and DNA damage repair by surviving cells. This hypothesis was confirmed when we analysed the nuclear localization of 53BP1 by quantitating *foci* positively stained with this marker (**Figure 5, c-e; Supplementary Figure 3**). The number of 53BP1-positive *foci* detected at 24hours post-irradiation increased proportionally to the priming concentration of ONC201 to which the cells have been exposed and was 6-fold greater with the highest doses (**Figure 5, c**). This led us to conclude that PC3 cells pre-treated with ONC201 were accumulating more DNA damage for a given dose of radiation, due to inhibitory effects on the expression of proteins required to resolve radiation-induced DNA damage and promote cell cycle re-entry. This pattern was maintained at a 72-hour timepoint post-irradiation, thus confirming that pre-treatment with ONC201 impaired the resolution of DNA damage post-irradiation (**Supplementary Figure 3, a**) (**Figure 5, d**).

**Figure 5.**
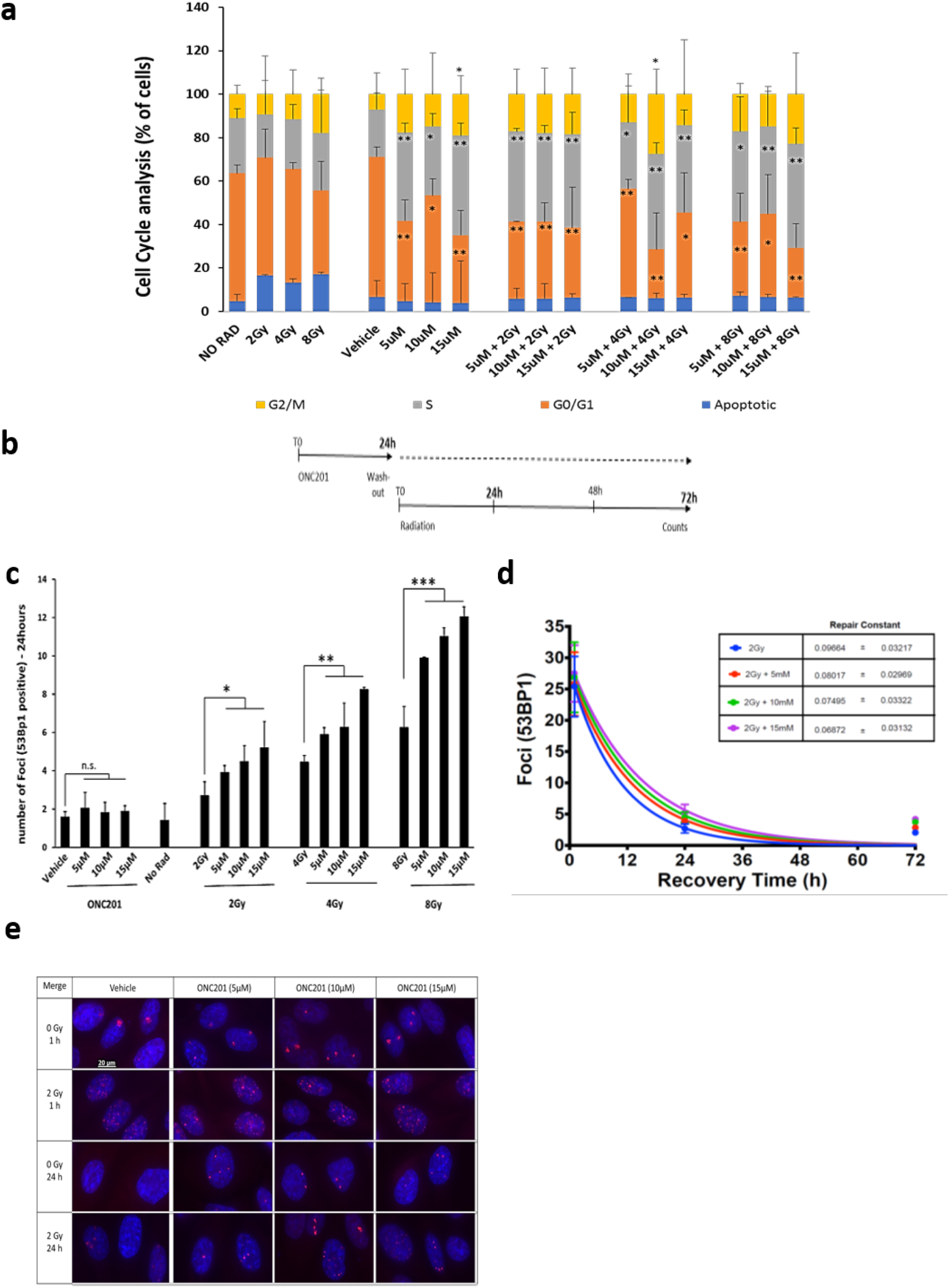
**a)** Cell Cycle analysis of PC3 primed to radiation (Xrad) with ONC201 for 24 hours shows an expansion of the cell population in S and G2/M phases. **b)** Schematic representation of the treatment in which PC3 cells have been primed with ONC201 before being radiated with single doses of Xrad. **c)** Number of 52BP1^+^ *foci* per cell at 24hours post irradiation. Samples were harvested as in scheme (b). **d)** Repair Kinetcs from the DNA damage induced by radiation. Samples were harvested as in scheme (b). **e)** FOCI determination through 53BP1 (red) staining at 1 and 24hours from radiation (IF). Samples were harvested as in scheme (b). One way ANOVA test has been run. The data shown in panel (c) have been run through ANOVA test on Ranks and further analysed with Dunnett’s Method. *P<0.05, **P<0.03, ***P <0.001 (±SD).

Based on these findings we hypothesized that drugs that could restrict cell cycle progression by directly inhibiting PLK1 or other cell cycle regulators might achieve similar sensitising effects if administered prior to radiation. We treated PC3 cells with the PLK1 inhibitor BI2536 (100nM) [35] (**Figure 6, a**). Pre-treating PC3 cells with this drug for 24 hours prior to radiation, in a similar manner to ONC201, led to a 50% enhancement in cell death. This is a more significant sensitization than we observed with the highest pre-treatment dose of ONC201. We also observed the same expansion that we observed with ONC201 pre-treatment in the proportion of cells in S phase and in the G2/M transition as well as increases in cell death by apoptosis and necrosis of around 20% (**Supplementary figure 4, b and c**). We also calculated the Radiation Enhancement Ratio (RER) to assess the impact of pre-treating cells with BI2536 on subsequent responses to radiation. At 72hours post-irradiation we increased the efficacy by three-fold when administering a 4Gy dose (**Figure 6, a**, right side).

**Figure 6.**
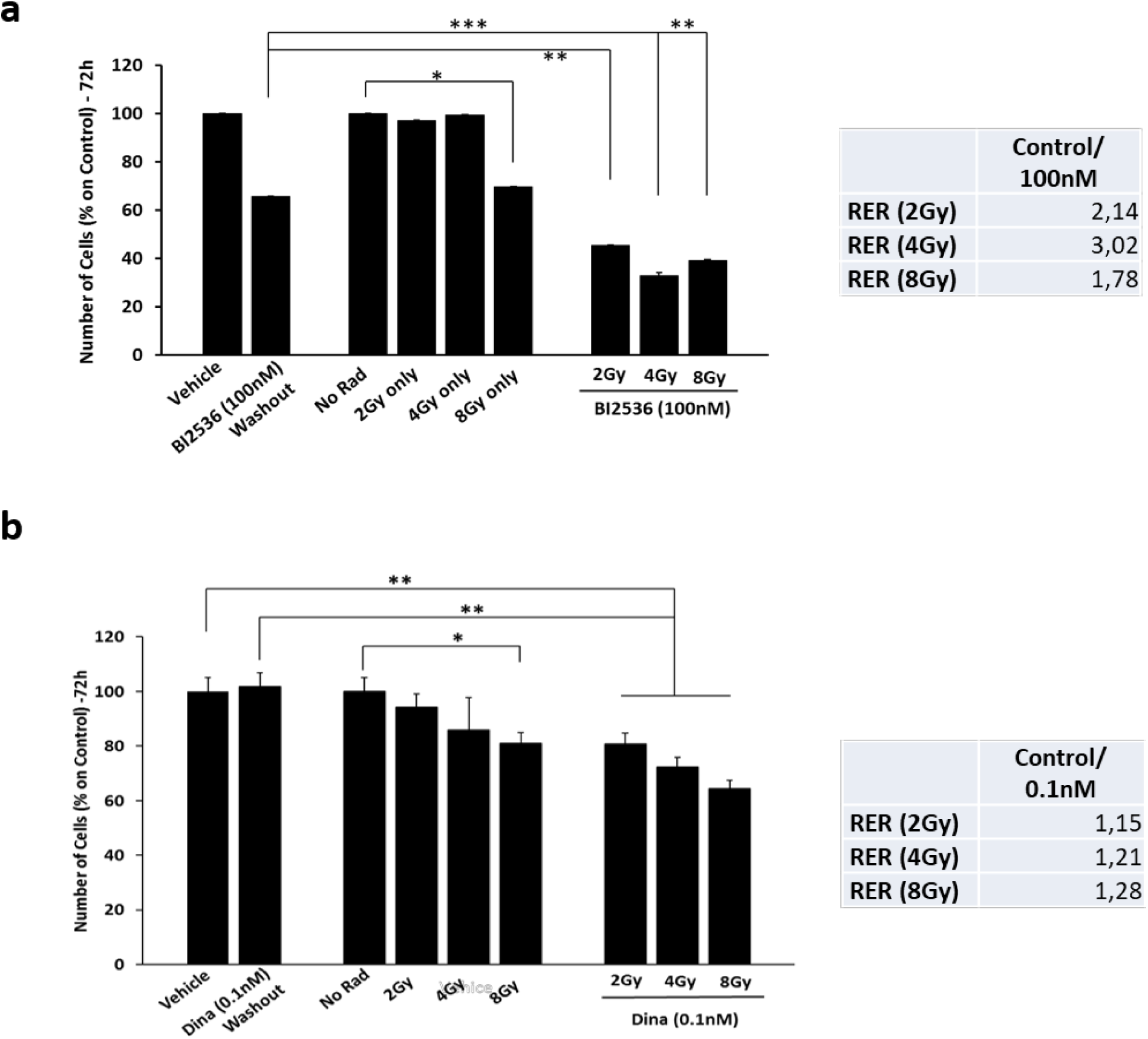
**a)** Cell counts of PC3 primed to radiation (Xrad) with the PLK1 inhibitor BI2536 (100nM) for 24hours. Counts in the plot show the number of cells at 72hours from radiation. Radiation Enhancement Ratio (RER) is shown and calculated out of 3 independent experiments. **b)** Cell counts of PC3 primed to radiation (Xrad) with a sub-toxic concentration of the CDKs inhibitor Dinaciclib for 24 (0.1nM). Radiation Enhancement Ratio (RER) is shown and calculated out of 3 independent experiments. One way ANOVA test has been run, *P<0.05, **P<0.03, ***P <0.001 (±SD).

Another cell cycle check point gene validated from the RNA-Seq analysis was CDK2 (**Figure 4, c and d**). CDK2 was downregulated by ONC201 at the mRNA level and upregulated by radiation alone (**Figure 4, d**). To assess whether this too could be exploited as a radiation sensitiser we pre-treated PC3 cells with a sub toxic dose of Dinaciclib, a CDK inhibitor (0.1nM) (**Figure 6, b**). By doing so, we were able to significantly reduce cell number by35% at 72hours and enhanced responses were once again confirmed by calculating the RER (**Figure 6, b**, right side). We also observed the same expansion in the proportion of cells in S phase and at the G2/M transition that we had observed with ONC201 pre-treatment as well as increased cell death by apoptosis and necrosis of ca. 20% (**Supplementary Fig.5, a and b**).

To determine whether the treatment sequence is important we subjected PC3 cells that had survived hypo-fractionated doses of radiation (5 times 2Gy and 10 times 2Gy) to ONC201 and found that prior exposure to radiation led to acquired resistance to ONC201 (**Supplementary figure 4, d and e**).

### ONC201 pre-treatment enhances radiation response in other cancer types

Given the sensitizing effects of ONC201 on PC3 prostate cancer cells to radiation (**Figure 2, a and b**), we wanted to assess whether ONC201 could have similar effects on other cell and cancer types. We therefore treated HT-29, a colorectal cancer cell line, with ONC201 (5, 10 and 15 µM) for 24hours and tested whether we were able to activate all the arms of the UPR (**Figure 7, a**). ONC201 at a concentration of 10µM increased the expression of proteins in all the arms of the UPR. We primed HT-29 cells to radiation with ONC201 (5 and 10 µM) for 24 hours. We followed up with 3 doses of radiation (2, 4 and 8Gy) (**Figure 7, b**). By doing so, we increased cell death by 50% (**Figure 7, b**) and enhanced apoptosis by 40% (**Figure 7, c**). The RER calculation confirmed this sensitisation effect showing once again a dose-dependent three-fold increase in efficacy when cells were pre-treated with ONC201 for 24 hours (**Figure 7, b**, right side). Together these data indicate that these radio-sensitisation effects may be broadly applicable and future studies will extend this work to additional models.

**Figure 7.**
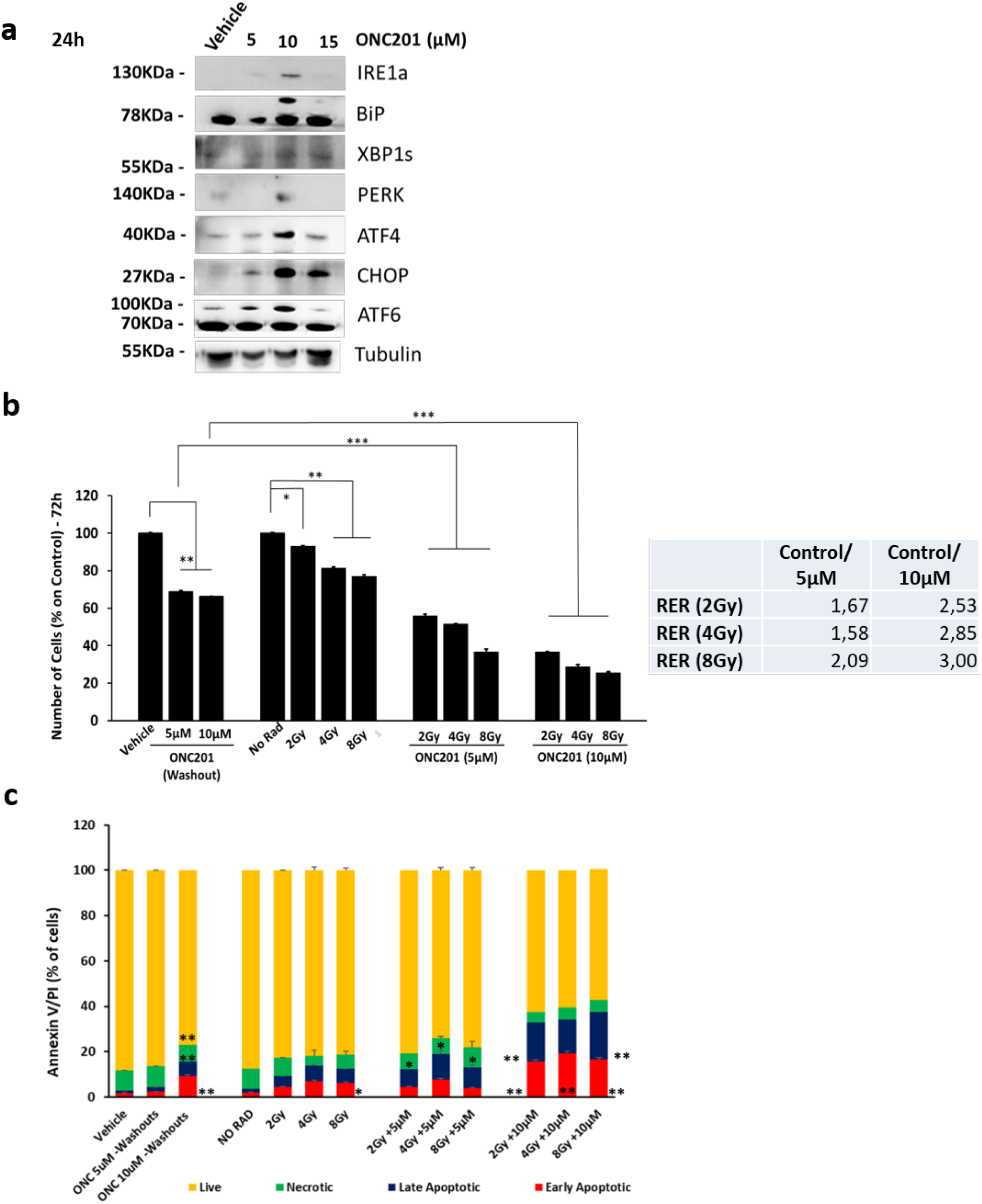
**a)** Treating colorectal cancer cells HT29 with ONC201 (5 and 10µM) upregulates all the proteins composing the UPR after 24hours. Representative of n=3 WB analysis of the expression of all the components of the UPR upon priming HT29 cells to radiation (Xrad) with ONC201 for 24hours. Densitometry analysis of n=3 WB are shown in Supplementary Fig.1, (b). **b)** Priming HT29 colorectal cells to radiation (Xrad) with ONC201 for 24hours impacts cells survival at 72hours. Radiation Enhancement Ratio (RER) is shown and calculated out of 3 independent experiments. **c)** Cell death analysis through Annaxin V/PI detection of PC3 treated with ONC201 (5 and 10µM) and Radiation (as described in a). Samples were harvested at 72h. One way ANOVA test has been run, *P<0.05, **P<0.03, ***P <0.001 (±SD).

## Discussion

Radiotherapy (brachytherapy, external beam radiation, and proton therapy) remains a highly effective modality in the treatment of multiple stages of prostate cancer. Between 10-15 % of patients are diagnosed after their cancer has spread and/or after first doses of radiation (with and without Hormone Castration –HDT-) fail, progressing to advanced or inoperable prostate cancer [4]. Understanding the molecular factors underpinning radiotherapy response and treatment failure is an area of research in which several signaling pathways having recently been implicated, including the UPR.

In this study we have assessed whether enhancing the expression of UPR components (particularly ATF4 and CHOP) in a sustained manner can synergise with radiation [37].

ATF4 activation can have both pro-apoptotic and pro-survival effects, in particular supporting cell-intrinsic metabolic adaptations [38-41]. In fact, tumour cells inside a growing tumour mass, often overexpress ATF4 to alleviate the stress from rapid proliferation and limited nutrient supply. The crucial impact of ATF4 activation on cell viability has been linked to the ATF4-dependent modulation of the balance between direct expression of adaptive versus pro-apoptotic targets and indirect control on targets involved in autophagy and protein synthesis [42]. Given the multifaceted activity of ATF4, we focused our attention on the downstream effector CHOP and through its targeted silencing we induced cell survival in prostate cancer (**Figure 1**). On the other hand, we expected that the activation of pro-apoptotic arms of the UPR may lead prostate cancer cells to cell death. To verify our hypothesis we tested a new compounds (ONC201) known to induce an integrated stress response ATF4- and CHOP-dependent in other cancer models and in phase I and II trials [2, 30](**Figure 2 and Supplementary figure 1, a**). For the first time, we observed an ONC201-induced activation of all the arms of the UPR in the first 24hours. ONC201 was then increasing cell death by apoptosis and necrosis at 72hours from the treatment. Although the canonical programmed cell death is known to be activated by unresolved UPR activation and autophagy, apoptosis has also been described through IRE1-dependent activation of JNK pro-apoptotic pathways [43]. Necrosis on the other hand has been linked to a sustained activation of the anticipatory UPR initiated by ATP depletion in ER^+^ breast cancer cells [44]. ONC201 induced necrosis in our hands but also induced an expansion of the S phase cell population. Increased expression of UPR components upon 24hours of ONC201 treatment, and prior to the induction of cell death, created a time window that we could exploit to assess whether short-term dosing could enhance radiation responses. Through pre-treatment we were able to increase the percentage of apoptotic and necrotic cells arising from radiation exposure by about 40% (**Figure 3, e**). ONC201 has previously been reported to modulate cell cycle progression in a model of haematological malignancies and in leukemic stem/progenitor cells where ONC201 causes p53-independent apoptosis and a delayed S phase transition [32]. In our hands we observed an S phase expansion as well as increased cell death. Sustaining cells in S phase and having them exposed to DNA damaging agents can lead to the accumulation of DNA aberrations and a concomitant build-up of misfolded/unfolded proteins in the endoplasmic reticulum. RNA-Seq analysis showed that when ONC201 was followed by radiation, this led to reduced expression of cell cycle and DNA damage repair pathway genes compared to radiation alone – RRM2, PLK1, CDK1, CDK2- (**Figure 4**). Radiotherapy is known to cause double-strand DNA damage [16]. The majority of these DNA double-strand breaks (DSBs) can be repaired by non-homologous end joining (NHEJ) through the whole cell cycle and by homologous recombination repair (HRR) during late S and G2 phases [45]. In both scenarios, histone alterations, nucleosome repositioning and changes in the higher-order folding of the chromatin fibre occur prior to the repair of lesions [46]. These modifications cause massive recruitment of the proteins gamma H2AX and 53Bp1 in large segments of the lesioned chromatin [47] [48] [49]. Because of these mechanisms, the S phase of the cell cycle is typically considered to be the phase that supports the emergence of radiation resistant cells. Collectively, our data suggest that cells persist in S phase through an inability (due to the mechanism of action of ONC201) to transition into M phase, making them more prone to the accumulation of unresolved radiation-induced DNA damage, ultimately resulting in cell death by necrosis or apoptosis.

To further test this hypothesis, we detected the accumulation of *foci* into the nuclei of cells primed to radiation with ONC201. Measuring the nuclear localization of γH2AX (data not shown) and 53Bp1 we were able to correlate the accumulation of DNA damage -and the delay in repairing it-to the phenotypical observations we had collected so far (**Figure 5, c-e**).

To further strengthen the evidence that ONC201 induces cell death in prostate cancer cells in part by impairing cell cycle progression, in addition to activating the ATF4/CHOP arm of the UPR, we tested two well-known inhibitors of PLK1 (BI2536) and CDK (Dinaciclib) (**Figure 6**). PLK1 and CDK1 are both regulate the G2/M DNA damage checkpoint. It ensures that cells don’t initiate mitosis until damaged DNA or incompletely replicated DNA is sufficiently repaired after replication (which occurs during the S phase, or Interphase) [50]. Cells that have a defective G2-M checkpoint enter mitosis before repairing their DNA, leading to apoptosis or death after cell division [50]. In our study, by treating prostate cancer cells with BI2536 or Dinaciclib for 24hours before radiation we were able to mimic the priming effect to radiation we obtained with ONC201 (**Figure 6**). We also increased apoptosis and necrosis as well as expanded the S phase of the Cell Cycle in both cases (**Supplementary figure 3 and Supplementary figure 4**). The use of cell cycle check points inhibitors in combination with radiation has already been proposed for the treatment of other cancer types such as Ewing’s sarcoma [51, 52], and is very consistent with the data we obtained with. Our study indicates for the first time that ONC201 may function as a radiation sensitiser in an analogous manner to other therapeutics that affect the cell cycle and are used (i.e. BI2536) and FDA approved (i.e. Dinaciclib) for the treatment of other cancer types (such as melanoma [53], CLL [54] and pancreatic cancer [55]). Although radiotherapy is the standard-of-care for several cancers (e.g. glioma, prostate cancer, etc.), in many cases is still not sufficient to eradicate the disease. Thus, combination of the molecule with radiotherapy will be the next step in designing clinical strategies and treatments. Understanding how treatments are used sequentially and how tumour cells adapt to therapies will also clearly be important. PC3 cells that survived hypo-fractionated doses of radiation (5 times 2Gy and 10 times 2Gy), were resistant to ONC201 (**Supplementary figure 4, d and e**). It is therefore possible that prior exposure to radiotherapy can promote the evolution of cells with altered responses to therapeutics affecting the UPR. Based on our current data we would propose further evaluation of the priming effect of ONC201 on radiotherapy-naïve prostate cancers. Future studies will however also need to account for a range of genomic backgrounds in those cancers and the duration and scheduling of such treatments as this work is expanded into a wider range of models and ultimately into a clinical setting.

## Material and Methods

### Proliferation and Annexin V/PI Assay

For the Proliferation assay, cells were treated as described in supplementary materials and maintained in culture up to 72hours from the administration of the treatment. When irradiated, for each experiment, unexposed controls were prepared and treated as sham exposures and harvested at matched time intervals. Adherent cells surviving radiation and/or drug administration where imaged in bright field (Leica CTR600, Leica, Wetzlar, Germany) at each time point and counted using ImageJ software (Public Domain, BSD-2). A minimum of 6 pics/well was taken as representative of the samples. Counts were normalized on corresponding T0. In the case of sequential experiments in which pre-treatment with ONC201, BI2536 and Dinaciclib was followed by irradiation, further analysis of the Survival Fractions (SF) -data not shown – and the Radiation Enhancement Ratio (RER) were performed as described in Subiel et al. [56].

For annexin/PI analysis, both floating and adherent cells were harvested and analysed with the FITC Annexin V Apoptosis Detection Kit I (BD Bioscience), according to the manufacturer’s instructions. Cells were analysed utilizing the FACS flow LSR II (BD Bioscience).

### Cell Cycle Analysis

Both floating and adherent cells were harvested at 72hours from the treatment (as described previously). After being washed in a solution of FBS in PBS (1:100), cells were fixed by adding ice cold 100% Ethanol to the solution of FBS-PBS (4:1). Afterwards, cells were stained by replacing the solution containing ethanol with another made up of PBS/FBS and implemented with PI (1:100, BD Bioscience) and RNaseA (1:400, Life Technologies). Cells were then analysed through FACS flow LSR II (BD Bioscience).

### Analysis of Synergism/Antagonism in combination treatments

The nature of the interaction between ONC201 and X-rays was determined using the CompuSyn software (ComboSyn, Inc. and [57]). For the purpose, the viability of cells after 72 hours from the treatments with ONC201 and radiation (alone and as sequential treatment) were inserted in the matrix of the Software to generate the algorithm.

### RNA-Seq Analysis

Cells were harvested at the time-points indicated in Figure 3, A. The RNA was extracted from the cells and collected utilizing the column-based method through the RNeasy MinElute Clean-up Kit (QIAGEN, Hilden, Germany), at a concentration of 25ng/µL in 20µL, according to the manufacturer’s instructions. The RNA library preparations were performed with the KAPA RNA HyperPrep Kit (KAPA Biosystems, Roche Holding AG) according to the manufacturer’s instructions. Sequencing was performed on Illumina Next Seq 500 (High out output (150)) at 75 P.E at 25M reads/ sample. This was performed by the Genomic Core Technology Unit, CCRCB, Queen’s University Belfast. Fastq files were generated using bcl2fastq version 2.19 using the default thresholds. Reads were aligned to version GRCh37/hg19 of the human reference genome using STAR (version 2.4.2a). Raw counts of reads mapped to genes were calculated using HT-Seq (http://www-huber.embl.de/users/anders/HTSeq/doc/overview.html) and used as input for differential expression analysis using DESeq2 [58] [59]. Principal components analysis was employed to visualise the overall effect of experimental covariates and batch effects within the data. Read counts per genes files were generate and principal component analysis (PCA) (RStudio, Boston, USA) performed. Gene sets were filtered to contain genes upregulated with radiation (ratio >1.5) and downregulated with ONC201 (ratio <0.5). Pathway analysis were performed on this gene list using Reactome (Reactome.org, ELIXIR) and Enrichr [60] [61].

### Clonogenic Assays

Clonogenic assays were performed as previously reported [62].

### *FOCI* detection

2*10^4^ cells were seeded on 13mm sterilized coverslips and treated as described previously. Cells were fixed at each time point (1hours, 24hours and 72hours form each treatment). DNA damage was detected using the immunofluorescence assay as previously described [17]. Briefly, cells were fixed in cold 50:50 solution of acetone and methanol and therefore permeabilized in 0.5% Trinton X-100 (Merks Chemicals, Darmstadt, Germany) in PBS for 20 min and then incubated for 60 min in blocking buffer (0.1% Triton X-100, 5% FBS in PBS). Anti-53BP1 rabbit monoclonal primary antibody (Novus Biological, Centennial, USA), diluted 1:5000 in blocking buffer was added to the cells. Samples were incubated at room temperature for 1hour. Secondary antibody (anti-rabbit Alexa Fluor 568, Life Technologies, Carlsbad, USA) diluted 1:2000 in blocking buffer was added and incubated for 1 hour. Cells on cover slips were briefly dried before mounted on slides with Prolong Gold Antifade with DAPI (Thermo Fisher, Waltham, USA). The number of 53Bp1 positive *foci* per cell were counted and imaged with the Nikon Eclipse Ti fluorescent microscope (Nikon Instruments Inc., Melville, NY, U.S.A.) in the entire nucleus. Experiments were repeated three times and at least 50 cells per repeat, treatment and time point were analysed.

### Data Analysis

All data are shown as mean ± S.D. Tests of significance were performed by one-way ANOVA test, using Sigma Plot software (Systat Software Inc, Evanston, IL, USA). To analyse the statistical significance of the number of *foci* detected as previously described, data were run through the ANOVA test on Ranks (for not normalised distributions) followed by the many-to-one comparisons (Dunnet’s Method). Also in this case, Sigma Plot software was utilised. Significant changes had p-values <0.05 (* p<0.05; **p<0.03; *** p<0.001).

## Acknowledgements

This work was supported by **Movember Prostate Cancer UK Centre of Excellence (CEO13_2-004)** to JOS and DJW and also by a grant from the Norwegian Research Council (230559) to IGM. IGM is also supported by the John Black Charitable Research Foundation. The authors acknowledge the Genomic Core Technology Unit, CCRCB, Queen’s University Belfast that performed the RNA-Seq Sequencing; the support of Fundação para a Ciência e Tecnologia (FCT-MCTES), Radiation Biology and Biophysics Doctoral Training Programme (RaBBiT, PD/00193/2012); and scholarships grants number SFRH/BD/114448/2016 to (FL).

## References

1. Storm M. S.X., Jildouw Arnoldussen Y. and Saatcioglu F., Prostate cancer and the unfolded protein response. Oncotarget, 2016. 7(33): p. 54051–54066.

2. Allen JE, K.C., Prabhu VV, Wagner J, Ishizawa J, Madhukar N, Lev A, Baumeister M, Zhou L, Lulla A, Stogniew M, Schalop L, Benes C, Kaufman HL, Pottorf RS, Nallaganchu BR, Olson GL, Al-Mulla F, Duvic M, Wu GS, Dicker DT, Talekar MK, Lim B, Elemento O, Oster W, Bertino J, Flaherty K, Wang ML, Borthakur G, Andreeff M, Stein M, El-Deiry WS., Discovery and clinical introduction of first-in-class imipridone ONC201. Oncotarget, 2016. 7(45): p. 74380–74392.

3. Smittenaar CR, P.K., Stewart K, Moitt N., Cancer Incidence and Mortality Projections in the UK Until 2035.. Brit J Cancer, 2016. 115(9): p. 1147–1155.

4. Nørgaard M1, J.A., Jacobsen JB, Cetin K, Fryzek JP, Sørensen HT., Skeletal related events, bone metastasis and survival of prostate cancer: a population based cohort study in Denmark (1999 to 2007). The Journal of urology,, 2010. 184(1): p. 162–7.

5. K., M., Tripartite Management of Unfolded Proteins in the Endoplasmic Reticulum. Cell, 2000. 101: p. 451–454.

6. Lu, P.D., Jousse, C., Marciniak, S. J., Zhang, Y., Novoa, I., Scheuner, D., Kaufman, R. J., Ron, D. and Harding, H. P., Cytoprotection by pre-emptive conditional phosphorylation of translation initiation factor 2. EMBO J., 2004. 23(169-79).

7. Lu, Y., Liang, F. X. and Wang, X., A synthetic biology approach identifies the mammalian UPR RNA ligase RtcB. Mol. Cell, 2014. 55: p. 758–770.

8. Lee, A.-H., Iwakoshi, N. N. and Glimcher, L. H., XBP-1 regulates a subset of endoplasmic reticulum resident chaperone genes in the unfolded protein response. Mol. Cell. Biol., 2003. 23: p. 7448–7459.

9. Kokame, K., Kato, H. and Miyata, T., Identification of ERSE-II, a new cis-acting element responsible for the ATF6-dependent mammalian unfolded protein response. J. Biol. Chem., 2001(276): p. 9199–9205.

10. Madden E, L.S., Healy SJ, Manie S, Samali A, The role of the unfolded protein response in cancer progression: From oncogenesis to chemoresistance. Biol Cell., 2019. 111 (1): p. 1–17.

11. Shuda M, K.N., Imazeki N, Tanaka K, Okada T, Mori K, Hada A, Arai M, Wakatsuki T, Matsubara O, Yamamoto N, Yamamoto M., Activation of the ATF6, XBP1 and grp78 genes in human hepatocellular carcinoma: a possible involvement of the ER stress pathway in hepatocarcinogenesis’. Journal of Hepatology, 2003. 38(5): p. 605–14.

12. Wärri A, C.K., Hu R, Jin L, Zwart A, Soto-Pantoja DR, Liu J, Finkel T, Clarke R., Autophagy and unfolded protein response (UPR) regulate mammary gland involution by restraining apoptosis-driven irreversible changes. Cell Death Discov., 2018. 4(40).

13. McCullough, K.D., Martindale, J. L., Klotz, L.-O., Aw, T.-Y. and Holbrook, N. J., Gadd153 sensitizes cells to endoplasmic reticulum stress by down-regulating Bcl2 and perturbing the cellular redox state. Mol. Cell. Biol., 2001. 21: p. 1249–1259.

14. Lu, M., Lawrence, D. A., Marsters, S., Acosta-Alvear, D., Kimmig, P., Mendez, A. S., Paton, A. W., Paton, J. C., Walter, P. and Ashkenazi, A., Opposing unfolded-protein-response signals converge on death receptor 5 to control apoptosis. Science 2014. 345: p. 98–101.

15. Decui Shao, J.N., Yang Shen, Jia Liu, Li Zhou, Hong Xue, Yu Huang, Wei Zhang, Limin Lu, CHOP mediatesXBP1S-inducedrenalmesangialcellnecrosisfollowing high glucosetreatment. Eur J Pharmacol., 2015. 758: p. 89–96.

16. Alice Sollazzo, B.B., Lei Cheng, Lovisa Lundholm, Harry Scherthan and a. AndrzejWojcik, Live Dynamics of 53BP1 Foci Following Simultaneous Induction of Clustered and Dispersed DNA Damage in U2OS Cells. International Journal of Molecular Sciences, 2018. 19(519).

17. Colman Trainor, K.T.B., Conor K. McGarry, Stephen J. McMahon, Joe M. O’Sullivan, Alan R. Hounsell, Kevin M. Prise, DNA Damage Responses following Exposure to Modulated Radiation Fields. PLoS One. 2012;, 2012. 7(8).

18. McLaughlin KA, N.Z., Bradley CA, Humphreys L, Stasik I, Fenning C, Majkut J, Higgins C, Crawford N, Holohan C, Johnston PG, Harrison T, Hanna GG, Butterworth KT, Prise KM, Longley DB., FLIP: A Targetable Mediator of Resistance to Radiation in Non-Small Cell Lung Cancer. Mol Cancer Ther, 2016. 15(10): p. 2432–2441.

19. Nagelkerke A, B.J., van der Kogel AJ, Sweep F C.G.J., Spana PN., The PERK/ATF4/LAMP3-arm of the unfolded protein response affects radioresistance by interfering with the DNA damage response. Radiotherapy and Oncology, 2013. 108(3): p. 415–421.

20. Shah SS., R.G., Musick A, Walters WM, de Cordoba N, Barbarite E, Marlow MM, Marples B, Prince JS, Komotar RJ, Vanni S and Graham RM, Targeting Glioblastoma Stem Cells with 2-Deoxy-D-Glucose (2-DG) Potentiates Radiation-Induced Unfolded Protein Response (UPR). Cancers, 2019. 11(159).

21. Niu, Z.W. M.; Zhou, L.; Yao, L.; Liao, Q.; Zhao, Y., Elevated GRP78 expression is associated with poor prognosis in patients with pancreatic cancer. Sci. Rep., 2015. 5(16067).

22. Nami B., G.-D.A., Vaseghi A., Overexpression of molecular chaperons GRP78 and GRP94 in CD44(hi)/CD24(lo) breast cancer stem cells. Bioimpacts, 2016. 6: p. 105–110.

23. Zhang J., J.Y., Jia Z., Li Q., Gong, W., Wang L., Wei D, Yao J., Fang S., Xie K., Association of elevated GRP78 expression with increased lymph node metastasis and poor prognosis in patients with gastric cancer. Clin. Exp. Metastasis, 2006. 23: p. 401–410.

24. Drake TM, R.J., Kanthou C, Staves JJ, Narramore R, Wyld L, Targeting the endoplasmic reticulum mediates radiation sensitivity in colorectal cancer. Exp Mol Pathol., 2015. 98(3): p. 532–9.

25. Avital Lev, A.R.L., Brian C. Ross, Marie D. Ralff, Peter B. Makhov, David T. Dicker and Wafik S. El-Deiry, ONC201 Targets AR and AR-V7 Signaling, Reduces PSA and Synergizes with Everolimus in Prostate Cancer. Mol Cancer Res., 2018. 16(5): p. 754–766.

26. Mark N. Stein, J.M., Rohinton S. Tarapore, Usha Malhotra, Ann W. Silk, Nancy Chan,Lorna Rodriguez, Joseph Aisner, Robert D. Aiken, Tina Mayer, Bruce G. Haffty, Jenna H. Newman,Salvatore M. Aspromonte, Praveen K. Bommareddy, Ricardo Estupinian, Charles B. Chesson, Evita T. Sadimin,Shengguo Li, Daniel J. Medina, Tracie Saunders, Melissa Frankel, Aparna Kareddula, Sherrie Damare,Elayne Wesolowsky, Christian Gabel, Wafik S. El-Deiry, Varun V. Prabhu, Joshua E. Allen, Martin Stogniew,Wolfgang Oster, Joseph R. Bertino, Steven K. Libutti, Janice M. Mehnert and Andrew Zloza., Safety and enhanced immunostimulatoryactivity of the DRD2 antagonist ONC201 in inadvanced solid tumor patients with weeklyoral administration. Journal for Immunotherapy of Cancer, 2019. 7(136).

27. Stein MN, B.J., Kaufman HL, Mayer T, Moss R, Silk A, Chan N, Malhotra J, Rodriguez L, Aisner J, Aiken RD, Haffty BG, DiPaola RS, Saunders T, Zloza A, Damare S, Beckett Y, Yu B, Najmi S, Gabel G, Dickerson S, Zheng L, El-Deiry WS, Allen JE, Stogniew M, Oster W, Mehnert JM., First-in-Human Clinical Trial of Oral ONC201 in Patients with Refractory Solid Tumors. Clin Cancer Res., 2017. 23(15): p. 4163–4169.

28. Sheng X, A.Y., Storm M, Tesikova M, Nenseth HZ, Zhao S, Fazli L, Rennie P, Risberg B, Wæhre H, Danielsen H, Mills IG, Jin Y, Hotamisligil G, Saatcioglu F., Divergent androgen regulation of unfolded protein response pathways drives prostate cancer. EMBO Mol Med., 2015 7(6): p. 788–801.

29. Lori S. Hart, J.T.C., Tatini Datta, Souvik Dey, Feven Tameire, Stacey L. Lehman, Bo Qiu, Haiyan Zhang, George Cerniglia, Meixia Bi, Yan Li, Yan Gao, Huayi Liu, Changhong Li, Amit Maity, Andrei Thomas-Tikhonenko, Alexander E. Perl, Albert Koong, Serge Y. Fuchs, J. Alan Diehl, Ian G. Mills, Davide Ruggero, and Constantinos Koumenis, 2012. J Clin Invest., ER stress– mediated autophagy promotes Myc-dependent transformation and tumor growth. 122(12): (12): p. 4621–4634.

30. Forsythe N, R.A., Javadi A, Khawaja H, Weir JA, Emam H, Allen WL, Burkamp F, Popovici V, Jithesh PV, Isella C, LaBonte MJ, Mills IG, Johnston PG, Van Schaeybroeck S., The unfolded protein response: a novel therapeutic target for poor prognostic BRAF mutant colorectal cancer. Mol Cancer Ther., 2018. pii: molcanther.0603.2017.

31. Greer YE1, P.-S.N., Nagashima K3, Stuelten C2, Crooks D4, Koparde VN5, Gilbert SF1, Islam C1, Ubaldini A1, Ji Y6, Gattinoni L6, Soheilian F3, Wang X7, Hafner M7, Shetty J8, Tran B8, Jailwala P5, Cam M5, Lang M4, Voeller D1, Reinhold WC9, Rajapakse V9, Pommier Y9, Weigert R2, Linehan WM4, Lipkowitz S.1, ONC201 kills breast cancer cells in vitro by targeting mitochondria. Oncotarget, 2018. 9(26): p. 18454–18479.

32. Jo Ishizawa, K.K., Dhruv Chachad, Peter P. Ruvolo, Vivian R Ruvolo, Rodrigo Jacamo, Archana Dilip, Hong Mu, Zhihong Zeng, Polina Matre, Joshua E. Allen, et al., ONC201 Induces p53-Independent Apoptosis and Cell Cycle Arrest in Hematological Malignancies and Leukemic Stem/Progenitor Cells By Inducing ER Stress and mTOR Inhibition. Blood, 2014. 124(3122).

33. Gülow K B.D.a.H.I., BiP is feed-back regulated by control of protein translation efficiency. Journal of Cell Science 2002. 115: p. 2443–2452.

34. Himanshi Bhatia, G.V., Malabika Datta, miR-107 orchestrates ER stress induction and lipid accumulation by post-transcriptional regulation of fatty acid synthase in hepatocytes. Biochimica et Biophysica Acta (BBA) – Gene Regulatory Mechanisms, 2014. 1839(4): p. 334–343.

35. Luying Jia, B.L.H.Y., The Bub1–Plk1 kinase complex promotes spindle checkpoint signalling through Cdc20 phosphorylation. Nature Communication., 2016. 7:10818.

36. Sharma NL, M.C., Ramos-Montoya A, Zecchini V, Scott HE, Lamb AD, MacArthur S, Stark R, Warren AY, Mills IG, Neal DE., The androgen receptor induces a distinct transcriptional program in castration-resistant prostate cancer in man. Cancer Cell, 2013. 23(1): p. 35–47.

37. Sano R, R.J., ER stress-induced cell death mechanisms. Biochim Biophys Acta., 2013. 1833(12): p. 3460–3470.

38. Hao Y, e.a., Oncogenic PIK3CA mutations reprogram glutamine metabolism in colorectal cancer. Nat Commun., 2016. 7(11971).

39. van Geldermalsen M, e.a., ASCT2/SLC1A5 controls glutamine uptake and tumour growth in triple-negative basal-like breast cancer. Oncogene, 2016. 35: p. 3201–3208.

40. Ren P, e.a., ATF4 and N-Myc coordinate glutamine metabolism in MYCN-amplified neuroblastoma cells through ASCT2 activation. J Pathol., 2015. 235(90-100).

41. Bi M, e.a., ER stress-regulated translation increases tolerance to extreme hypoxia and promotes tumor growth. EMBO J., 2005. 24: p. 3470–3481.

42. Wortel INN, v.d.M.L., Kilberg MS, and van Leeuwen FN., Surviving Stress: Modulation of ATF4-Mediated Stress Responses in Normal and Malignant Cells. Trends Endocrinol Metab., 2017. 28(11): p. 794–806.

43. Merksamer, P.I.P. F. R., The UPR and cell fate at a glance. J. Cell Sci., 2010. 123: p. 1003–1006.

44. Livezey M, H.R., Hergenrother PJ, Shapiro DJ., Strong and sustained activation of the anticipatory unfolded protein response induces necrotic cell death. Cell Death Differ., 2018. 10: p. 1796–1807.

45. Branzei D, F.M., Regulation of DNA repair throughout the cell cycle. Nat Rev Mol Cell Biol., 2008. 9(4): p. 297–308.

46. Panier, S.B. S.J., Double-strand break repair: 53BP1 comes into focus. Nat. Rev. Mol. Cell Biol., 2014. 15: p. 7–18.

47. Lukas, J.L. C.; Bartek, J., More than just a focus: The chromatin response to DNA damage and its role in genome integrity maintenance. Nat. Cell Biol., 2011. 13: p. 1161–1169.

48. Rogakou, E.P.B. C.; Redon, C.;, Bonner, W.M. Megabase chromatin domains involved in DNA double-strand breaks in vivo. J. Cell Biol., 1999. 146: p. 905–916.

49. Lottersberger, F.K. R.A.; Dimitrova, N.; de Lange, T. 53BP1 and the LINC Complex Promote Microtubule-Dependent DSB Mobility and DNA Repair. Cell 2015. 163: p. 880–893.

50. Cuddihy, A.R.O.C. M. J., Cell-cycle responses to DNA damage in G2. International Review of Cytology, 2003(222): p. 99–140.

51. K., P.T.a.K., Role of cell cycle in mediating sensitivity to radiotherapy. Int. Journal of Rad. Oncol., 2004. 59(4): p. 928–942.

52. Maier P, H.L., Wenz F and Herskind C., Cellular Pathways in Response to Ionizing Radiation and Their Targetability for Tumor Radiosensitization. Int. J. Mol. Sci., 2016. 17(102).

53. Desai BM, V.J., Nguyen TT, Lioni M, Xiao M, Kong J, Krepler C, Vultur A, Flaherty KT, Nathanson KL, Smalley KS, Herlyn M, The anti-melanoma activity of dinaciclib, a cyclin-dependent kinase inhibitor, is dependent on p53 signaling. PLOS ONE, 2013. 8(3).

54. Johnson AJ, Y.Y., Smith LL, Wagner AJ, Hessler J, Gupta S, Flynn J, Jones J, Zhang X, Bannerji R, Grever MR, Byrd JC The novel cyclin-dependent kinase inhibitor dinaciclib (SCH727965) promotes apoptosis and abrogates microenvironmental cytokine protection in chronic lymphocytic leukemia cells. Leukemia, 2012. 26(12): p. 2554–7.

55. Feldman G, M.A., Bisht S, Karikari C, Garrido-Laguna I, Rasheed Z, Ottenhof NA, Dadon T, Alvarez H, Fendrich V, Rajeshkumar NV, Matsui W, Brossart P, Hidalgo M, Bannerji R, Maitra A, Nelkin BD Cyclin-dependent kinase inhibitor Dinaciclib (SCH727965) inhibits pancreatic cancer growth and progression in murine xenograft models. Cancer Biology & Therapy, 2011. 12(7): p. 598–609.

56. Subiel A, A.R., Schettino G., Standards and Methodologies for Characterizing Radiobiological Impact of High-Z Nanoparticles. Theranostics, 2016. 6(10): p. 1651–1671.

57. Tc, C., Drug combination studies and their synergy quantification using the Chou-Talalay method. Cancer Res., 2010. 70: p. 440–446.

58. Dobin A, D.C., Schlesinger F, Drenkow J, Zaleski C, Jha S, et al., STAR: Ultrafast universal RNA-seq aligner. Bioinformatics, 2013. 29: p. 15–21.

59. Love MI, H.W., Anders S., Moderated estimation of fold change and dispersion for RNA-seq data with DESeq2. Genome Biol., 2014. 15(550).

60. Chen EY, T.C., Kou Y, Duan Q, Wang Z, Meirelles GV, Clark NR, Ma’ayan A., Enrichr: interactive and collaborative HTML5 gene list enrichment analysis tool. BMC Bioinformatics, 2013. 128(14).

61. Kuleshov MV, J.M., Rouillard AD, Fernandez NF, Duan Q, Wang Z, Koplev S, Jenkins SL, Jagodnik KM, Lachmann A, McDermott MG, Monteiro CD, Gundersen GW, Ma’ayan A., Enrichr: a comprehensive gene set enrichment analysis web server 2016 update. Nucleic Acids Research, 2016.

62. Butterworth KT, M.C., O’Sullivan JM, Hounsell AR, Prise KM., A study of the biological effects of modulated 6 MV radiation fields. Phys Med Biol., 2010. 55(1607-1618).

