## Supplementary materials and methods, supplementary figures 1-5 and supplementary tables 1 and 2 for "Modulating the unfolded protein response: Impacts of radiation on the response of prostate cancer cells to ONC201"

### Supplementary - Material and Methods

**Cell lines and reagents.** Prostate adenocarcinoma PC3 cell line and colorectal cancer HT-29 cell line were obtained from the American Type Culture Collection (Authentication by short tandem repeat profiling/karyotyping/isoenzyme analysis). PC3 cells were maintained in RPMI-1640 (GIBCO-Life Technologies, Thermo Scientific, UK) implemented with 10% FBS (GIBCO-Life Technologies), 2 mM L-Glutamine, 100 U/ml penicillin and 100 mg/ml streptomycin (GIBCO-Life Technologies). HT-29 cells were maintained in DMEM (GIBCO-Life Technologies, Thermo Scientific, UK) implemented with 10% FBS (GIBCO-Life Technologies), 2 mM L-Glutamine, 100 U/ml penicillin and 100 mg/ml streptomycin (GIBCO-Life Technologies). Cells were maintained at 37°C in a 5% $CO_2$-humidified atmosphere. All experiments were performed in complete medium. Cells were plated overnight at a known density. The medium was replaced and implemented with several different treatments (either drugs or radiation) up to the next 72 hours. When cells were irradiated medium was replaced immediately prior to each irradiation dose using an XRad 225 cabinet source at dose rate 0.59 Gy/min, (Precision X-ray Inc, North Branford, CT, USA). Unexposed controls were prepared and treated as sham exposures. ONC201 has been kindly provided by Oncoceutics (Philadelphia, PA, USA). ONC201 was dissolved as a stock solution of 20 mM in DMSO and used for experiments at a final concentration of 5, 10 and 15 µM in complete medium. HA15 (kindly supplied by Dr. S. Van Schaeybroeck, Queen's University Belfast) has been dissolved as a stock solution of 10 mM in DMSO and used for experiments at a final concentration of 1, 2.5 and 5 µM in complete medium [56]. BI2536 was purchased by Selleckchem (Munich, Germany). It has been dissolved as a stock solution of 10 mM in DMSO and used for experiments at a final concentration of 100 nM in complete medium [57]. The Vehicle was in all cases DMSO 0.1% in complete medium. The vector containing the short-hairpin RNA (shRNA) targeting DDIT3 (CHOP) -with sequence GGAAAGTGGCACAGCTAGC - belongs to a shRNA retroviral library generated at the Netherlands Cancer Institute (NKI) and described by

Bernards et al. [58]. The library is at our disposal in the Centre (CCRCB, Queen's University Belfast). The empty vector used to generate the Scram cell line is the pRSC vector (according to Bernards at al.) and has been purchased from Invitrogen (Life Technologies).

**Retroviral Infection.** Retrovirus generation was conducted as previously described [59].

**Western Blots.** Cells were treated with ONC201 alone for 24hours (Washouts) and up to 72hours, radiation alone (single doses of 2Gy, 4Gy and 8Gy) and as a sequential treatment with ONC201 for 24hours followed by radiation in single dose (a schematic view of the treatments is in Figure 3, A). Cells were detached trough trypsinization (5 minutes at 27˚C, Sigma Aldrich) at 24hours and 72hours after each treatment and lysed with Urea buffer (Urea 9 M, Tris Base 1 M, PMSF 10 mM, beta-mercaptoethanol 1%, (all Sigma Aldrich). Proteins were separated on 4-12% NuPage precast gels (Life Technologies) and blotted on nitrocellulose paper (Life Technologies). Membranes were incubated overnight with antibodies anti- IRE1a (14C10), BiP (C50B12), PERK (D11A8), CHOP (L63F7), PLK1 (208G4) (all Cell Signaling, Danvers, MA, USA), ATF4 (EPR18111) (Abcam, Cambridge, UK), ATF6 (70B1413.1) (Novus, Manchester, UK), PARP (46D11) and Caspase 3 (all detecting both full length and cleaved forms, Cell Signaling) at a dilution of 1:1000 in a solution of 3% not fat milk-PBS. Antibody anti-XBP1 spliced (143F) (Biolegend, San Diego, CA, USA), was used at a dilution of 1:400 in 3% not fat milk-PBS.

The anti-tubulin Ab (Cell Signaling) was used as housekeeping control at a dilution of 1:1000 as described above. The secondary anti-rabbit and anti-mouse horseradish peroxidase-conjugated Abs (Cell Signaling) were used at a dilution of 1:3000 in PBS-0.25% Tween (Sigma Aldrich). Protein bands were revealed with the Western Lightning Ultra kit (PerkinElmer, Waltham, MA, USA) using the GBox Imager by Syngene (Cambridge, UK). Densitometic analysis of the blots (n=3) were performed utilizing ImageJ software (Public Domain, BSD-2).

**qRT-PCR and primers.** To perform the RNA extraction, cells were treated and harvested at each time point trough trypsinization (as described above) and processed with Trizol (Life Technologies) according to the manufacturer's instructions.

qRT-PCR analysis was performed using the LightCycler® 480 probes master mix and LightCycler®480II (both Roche, Basel, Switzerland).

The expression levels of RPLPO gene have been used as housekeeping. The primers (all from Eurofin Genomics UK) used in this work have the following sequences:

| targeted gene (protein) | Forward sequence (5'-3') | Reverse Sequence (5'-3') |
|---|---|---|
| DDIT3 (CHOP) | cagatcaaaatgggggtaccta | tcaagagtggtgaagatttttgat |
| RRM2 | Tggacctctccaaggacatt | ggctaaatcgctccacca |
| ZBTB20 | atgggtgcacacaggaaaat | cttctcatgggcctgtatgtt |
| MKI67 (Ki67) | gagagtaacgcggagtgtca | tcactgtccctatgacttctgg |
| TYMS | Cccagtttatggcttccagt | gcagttggtcaactccctgt |
| PLK1 | aagatctggaggtgaaaataggg | aggagtcccacacagggtct |
| CDK1 | tggatctgaagaaatacttggattcta | tctggagatctgtaccagagtgtt |
| CDK2 | Cctcctgggctgcaaata | cagaatctccagggaataggg |
| RPLPO | atcaacgggtacaaacgagtc | cagatggatcagccaagaagg |

**Supplementary - Figure Legends**

**Supplementary Figure 1. a)** HA15 has a cytostatic effect on PC3 cells treated up to 72h. **b)** Priming PC3 cells to radiation with ONC201 for 24hours prevents colony formation. **c)** Representative of n=3 WB analysis of the expression of all the components of the UPR upon priming PC3 cells to radiation (Xrad) with ONC201 for 24hours reveals the activation of pro-apoptotic arms of the UPR (ATF4/CHOP). Samples were harvested at 24hours (left panel) from the last radiation. Samples treated with ONC201 for only 24hours and harvested at 72hours from the washouts (ONC201 W.O.) are also shown (right panel). **d)** Densitometry of n=3 WB are shown. One way ANOVA test has been run, *P<0.05, **P<0.03, ***P <0.001 (±SD).

**Supplementary Figure 2. a)** Densitometry of WB shown in Figures 3. The expression of all the components of the UPR upon priming PC3 cells to radiation (Xrad) with ONC201 for 72hours were analysed. Samples were harvested at 72hours from the last radiation. One way ANOVA test has been run, *P<0.05, **P<0.03, ***P <0.001 (±SD). **b)** PCA plots of RNAseq analysis of PC3 primed to radiation (Xrad) with ONC201 for 24 hours show two subgroups of differentially expressed gens (samples harvested at 24 hours from radiation –red dots- and at 72 hours – blue dots). One way ANOVA test has been run, *P<0.05, **P<0.03, ***P <0.001 (±SD). **c)** Pathway analysis reveals that the most impacted pathway is cell cycle related (http://cpdb.molgen.mpg.de/ - p-value cut-off = 0.01 with minimum overlap with input list = 2).

**Supplementary Figure 3. a)** Number of 52BP1$^{+}$ *foci* per cell at 24hours post irradiation. Samples were harvested as in Figure 5, (b). One way ANOVA test has been run. The data shown in panel C have been run through ANOVA test on Ranks and further analysed with Dunnett's Method. *P<0.05, **P<0.03, ***P <0.001 (±SD). **b)** Single channels of the *foci* determination at 1 and 24hours from radiation (IF). 53BPp1 (red) and DAPI (blue) are shown. Samples were harvested as in Figure 5, (b).

**Supplementary Figure 4. a)** WB analysis of the expression of PLK1, one of the major check points of the S/$G_2$M transition through the Cell Cycle. Samples were harvested at 72h from the last radiation. **b)** Cell cycle analysis at 72hours of PC3 treated with BI2536 (100nM) and Radiation. **c)** Inhibiting PLK1 with BI2536 (100nM) before radiation increases cell death through Necrosis and Apoptosis at 72hours. **d-e)** PC3 cells surviving hypo-fractionated doses of radiation (d, 5 times 2Gy – e, 10 times 2Gy respectively) are more resistant to ONC201 (5µM and 10µM). One way ANOVA test has been run, *P<0.05, **P<0.03, ***P <0.001 (±SD).

**Supplementary Figure 5. a)** Cell cycle analysis of PC3 treated with CDKs inhibitors at 72h. **b)** Cell death analysis of PC3 treated with CDKs inhibitors at 72h. One way ANOVA test has been run, *P<0.05, **P<0.03, ***P <0.001 (±SD).

**Table1.** Table with a list of the most differentially expressed genes shortlisted from the RNA-Seq analysis (Ratio>1.5) in relation to their direct regulation of the most effected pathways (Enrichr). The Venn diagram graphically represents this grouping (Venny 2.1.0 [63]).

**Table2.** Table with a list of the top three most impacted pathways by the most differentially expressed genes obtained through RNA-Seq analysis (Ratio>1.5), according to the Reactome tool.

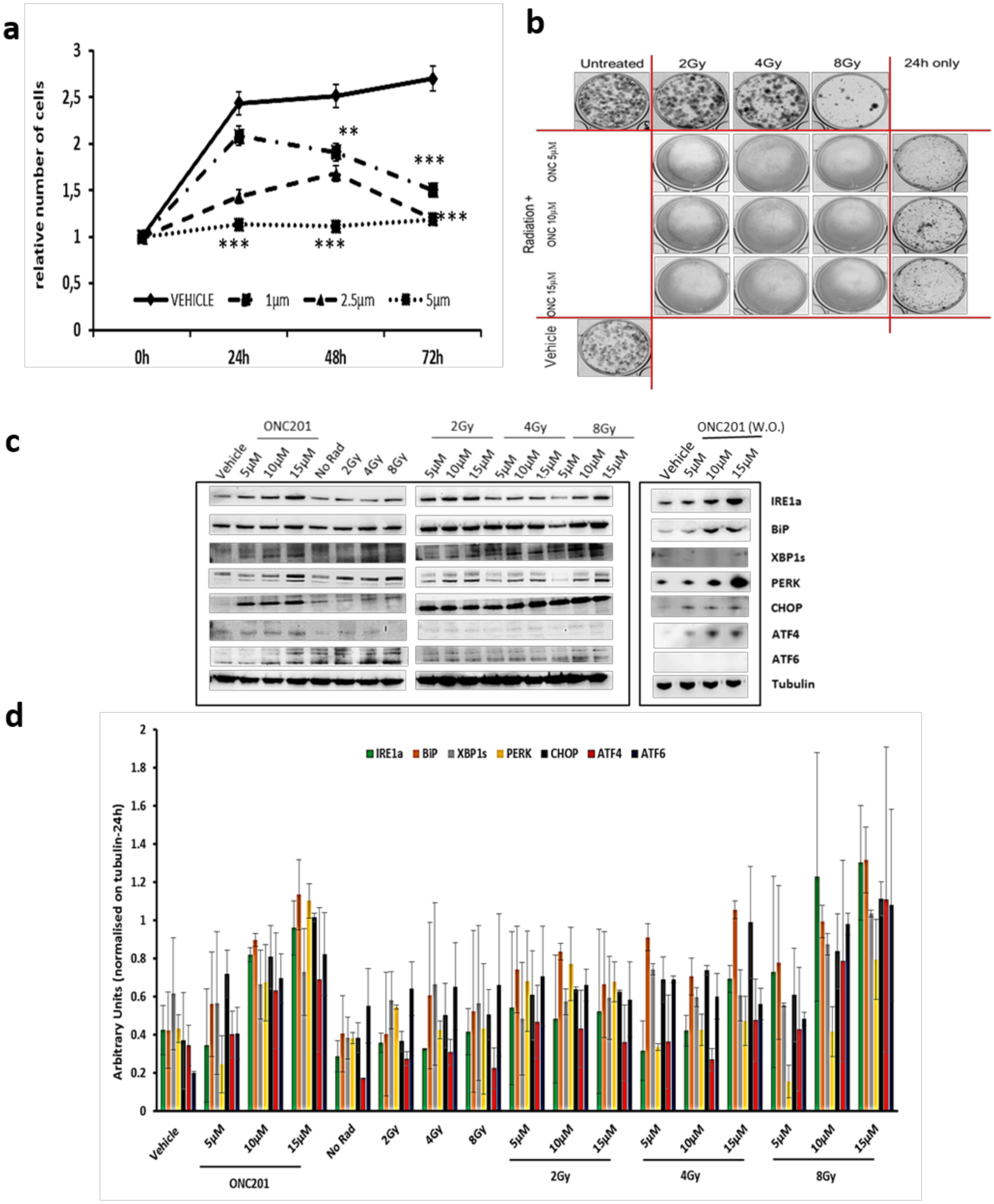


**Supplementary Figure 1.**

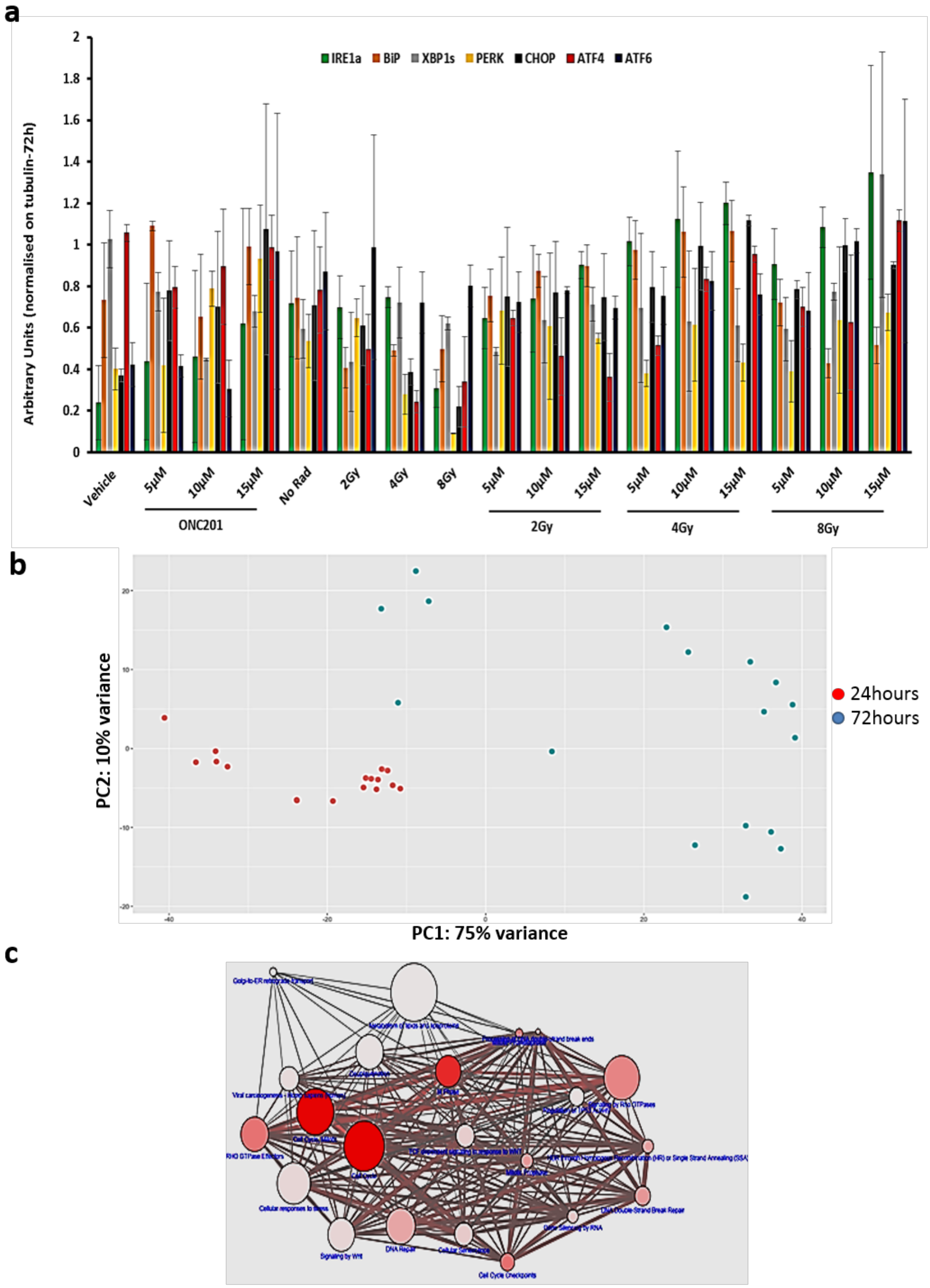


**Supplementary Figure 2.**

**a**

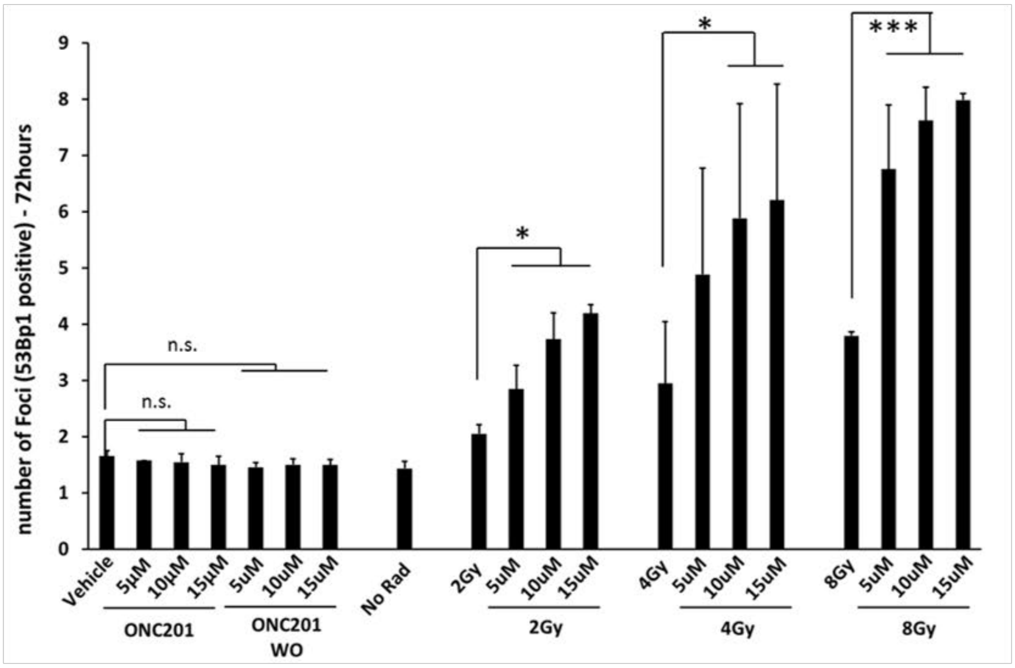


**b**

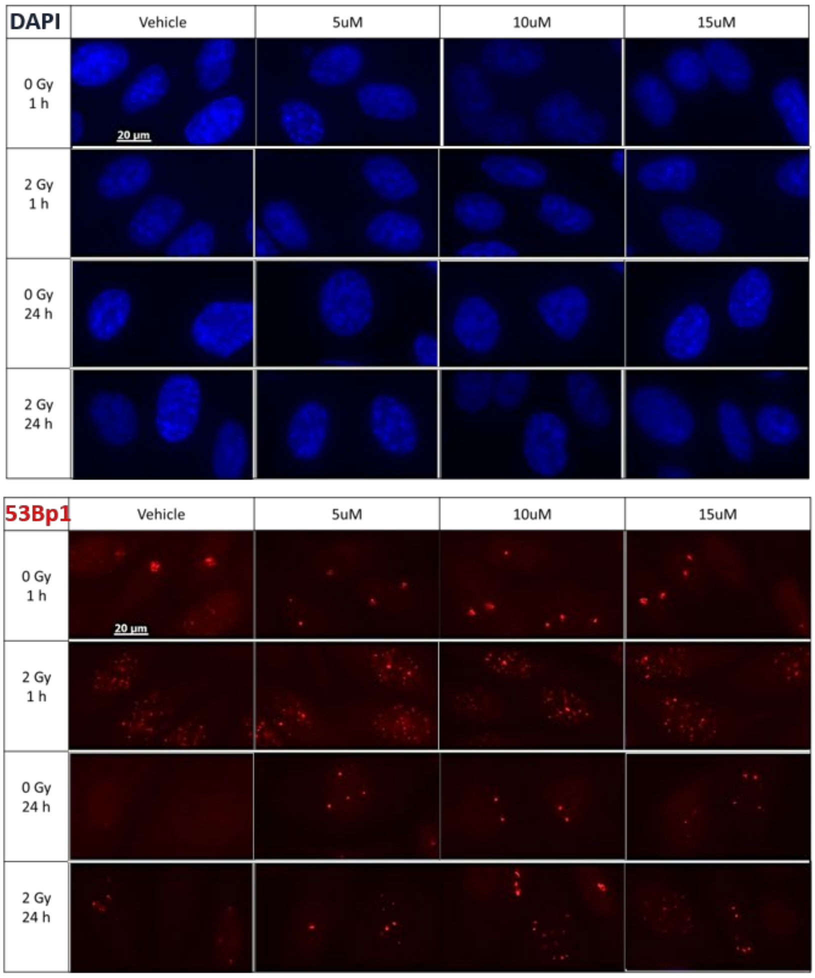


**Supplementary Figure 3.**

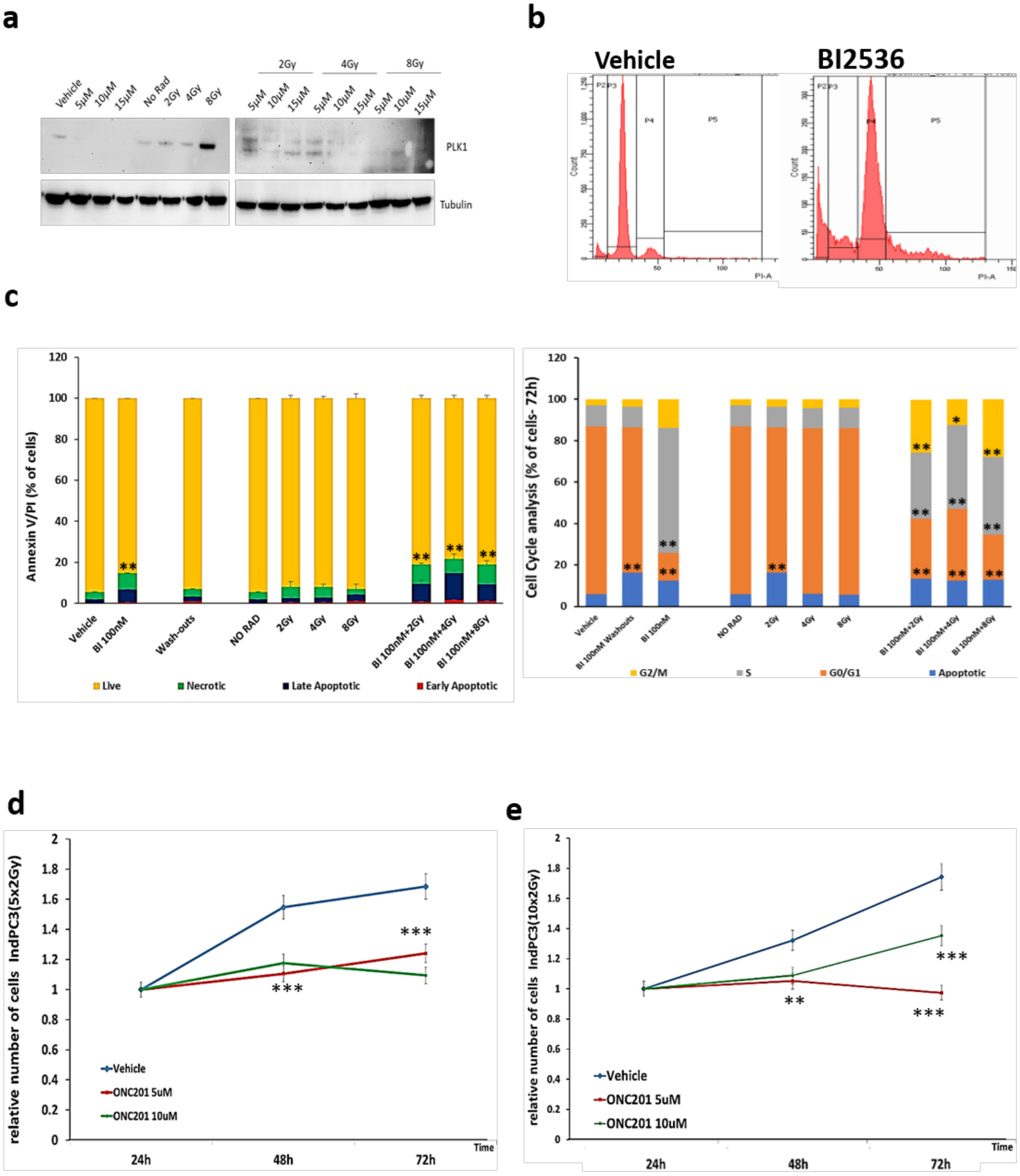


**Supplementary Figure 4.**

a

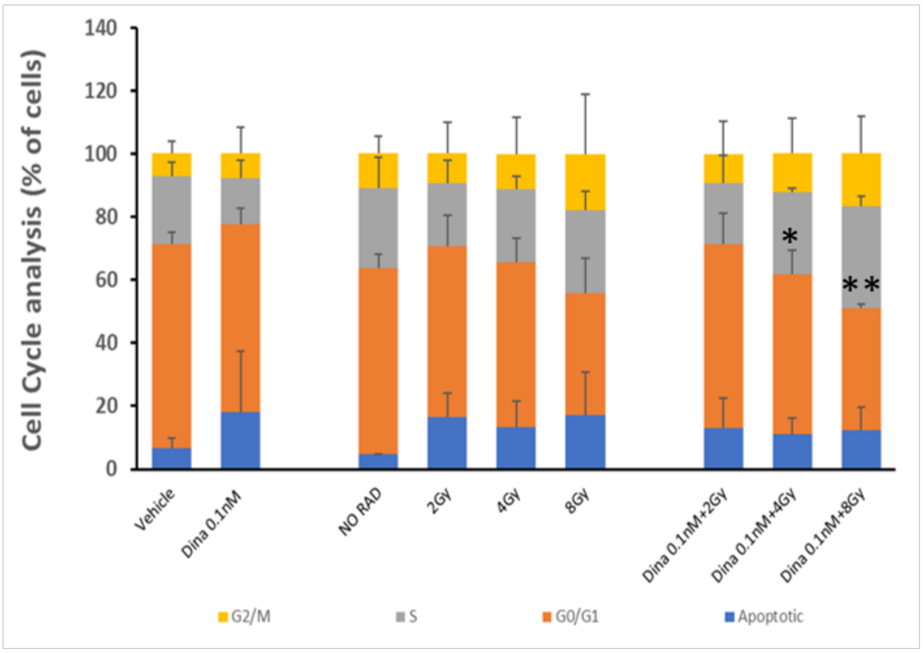


b

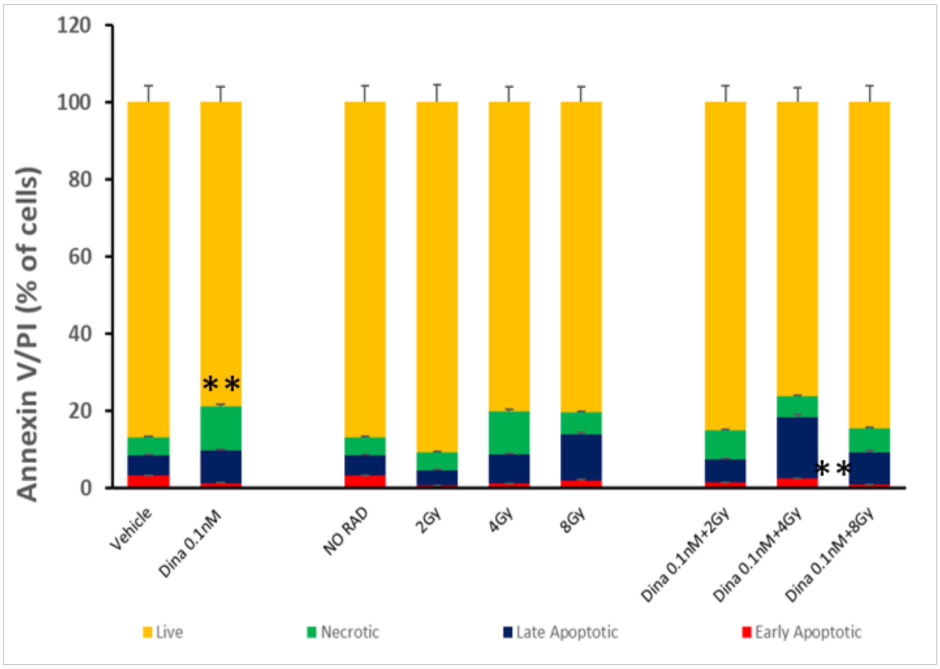


**Supplementary Figure 5.**

| common elements in "Ratio >1.5 Rad" and "Reactome_Cell Cycle" | common elements in "Ratio >1.5 Rad", "Reactome_Cell Cycle" and "Reactome_RHO effectors" | common elements in "Ratio >1.5 Rad", "Reactome_Cell Cycle" and "Reactome_DNArepair" | common elements in "Ratio >1.5 Rad" and "Reactome_RHO effectors" | common elements in "Ratio >1.5 Rad" and "Reactome_DNArepair" |
|---|---|---|---|---|
| CDK1 | PLK1 | FEN1 | PRC1 | XRCC3 |
| TYMS | INCENP | PCNA | KIF14 | KPNA2 |
| RRM2 | MAD2L1 | RFC3 | CIT | CLSPN |
| LMNB1 | CDCA8 | CDK2 | | DTL |
| KIF20A | KIF2C | | | FANCI |
| CDCA5 | SPC24 | | | FANCD2 |
| MYBL2 | BIRC5 | | | NEIL2 |
| SKP2 | SGOL2 | | | POLH |
| AURKA | CASC5 | | | POLM |
| KIF23 | NDC80 | | | TIMELESS |
| UBE2C | BUB1 | | | POLQ |
| NCAPD2 | BUB1B | | | TDP1 |
| NUP210 | DSN1 | | | RAD18 |
| HAUS8 | CENPE | | | |
| TOP2A | CENPC | | | |
| CCNB2 | CENPF | | | |
| CCNB1 | CDC20 | | | |
| MCM8 | ZWINT | | | |
| MCM4 | NUP37 | | | |
| TMPO | KNTC1 | | | |
| TPX2 | | | | |
| NCAPH | | | | |
| PTTG1 | | | | |
| LPIN1 | | | | |
| NCAPG | | | | |
| POLA2 | | | | |
| NCAPD3 | | | | |
| DBF4 | | | | |
| SMC2 | | | | |
| MCM6 | | | | |
| NCAPG2 | | | | |
| TUBB4B | | | | |

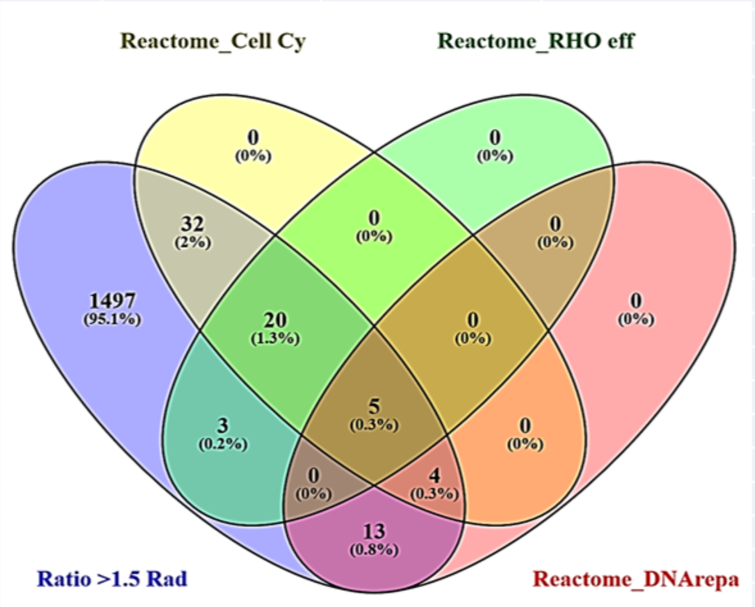


**Table1.**

| Top 3 pathways (Reactome) | | | |
|---|---|---|---|
| Index | bcl | Overlap | Adjusted p-value |
| 1 | Cell Cycle | 61/462 | 6.80977E-48 |
| 2 | RHO GTPase Effectors | 28/255 | 4.25709E-19 |
| 3 | DNA Repair | 22/285 | 6.6974E-12 |

**Table2.**
